# TMEM33 deletion potentiates anti-tumor CD8^+^ T cell immunity

**DOI:** 10.64898/2026.01.01.697288

**Authors:** Matthew T Jackson, Tianming Zhao, Gusztav Milotay, Isabela Pedroza-Pacheco, Yanshu Cai, Claire Willis, Su M Phyu, Bruno Beernaert, Jamie TW Kwon, Hala Estephan, Vinnycius Pereira Almeida, Maria Aggelakopoulou, Silvia Panetti, Christian Zierhut, Tim Elliott, Eleni Adamopoulou, Jan Rehwinkel, Malcolm JW Sim, Amit Grover, Dmitry I Gabrilovich, Benjamin P Fairfax, Eric Honoré, David R Withers, John C Christianson, Eileen E Parkes

## Abstract

Improving responses to cancer immunotherapies requires deeper insight into the cellular mechanisms governing T cell-mediated anti-tumor immunity. TMEM33 is an endoplasmic reticulum-resident transmembrane protein enriched across multiple tumor types, with reported functions in anti-viral immunity as well as calcium and lipid homeostasis, yet its role in tumor immunosurveillance remains unknown. Using murine genetic models, we demonstrate that host TMEM33 constrains anti-tumor CD8^+^ T cell responses. Constitutive *Tmem33^-/-^* mice exhibited delayed melanoma tumor growth and increased CD8^+^ T cell infiltration. Antigen-specific CD8^+^ compartments in tumors of *Tmem33^-/-^* mice showed TCF-1^+^PD-1^+^ progenitor-exhausted cell (Tpex) enrichment, elevated effector function and reduced exhaustion, alongside improved effector memory expansion and T-bet expression in draining lymph nodes. We highlight that TMEM33 functions intrinsically within the T cell compartment, as TMEM33 deletion (1) enhanced polyclonal activation of naive CD8^+^ T cells *ex vivo*, (2) promoted preferential Tpex accumulation among adoptively transferred naive OT-I cells in B16F10-OVA tumors and draining lymph nodes, and (3) improved the potency of *ex vivo*-expanded OT-I cells in controlling tumor growth during adoptive cell therapy. Finally, in a large, prospectively recruited metastatic melanoma cohort, lower *TMEM33* expression in patient CD8^+^ T cells significantly correlated with improved survival and elevated *TCF-7* (encoding TCF-1). Collectively, our findings define TMEM33 as a formerly unrecognized intrinsic determinant of tumor-directed CD8^+^ T cell fate that limits Tpex maintenance, and restrains cell therapy responses, suggesting that its modulation may strengthen immunotherapeutic efficacy.

**One sentence summary:** TMEM33 intrinsically limits progenitor exhausted CD8^+^ T cells, scales anti-tumor responses and predicts melanoma patient survival.

## INTRODUCTION

Despite the success of current immunotherapies in cancer treatment, the population of benefit remains limited (*1*). Deeper understanding of the cellular biology governing CD8^+^ T cell-mediated tumor control could explain response heterogeneity, and reveal novel, targetable mechanisms to improve clinical outcomes. While inhibitory and costimulatory molecules have largely been the focus of attention, additional fundamental cellular mechanisms that govern immune cell function in the tumor represent a critical yet unexplored layer of immune control. Many of these processes converge on the endoplasmic reticulum (ER) - a dynamic organelle central to tumor immunosurveillance, and include major histocompatibility complex (MHC) assembly (*2*), calcium (Ca^2+^) flux during lymphocytic activation (*3*), ER stress responses and innate immune signaling (*4*, *5*). These functions are scaled by an abundance of ER-resident regulatory proteins (e.g. E3 ubiquitin ligases) and functional cofactors (*6*, *7*), which collectively represent potential new avenues for immunotherapeutic intervention.

Transmembrane protein 33 (TMEM33) is an evolutionarily conserved, ubiquitously expressed protein positioned at the ER, where it acts as a structural cofactor and modulates the abundance and function of numerous binding partners. We previously identified TMEM33 as a component of an ER-embedded immunomodulatory complex, centered on the E3 ligase RNF26, where it attenuates type I interferon responses mediated by the innate immune signaling hub, STimulator of INterferon Genes (STING) (*8*). In cancer, TMEM33 expression is elevated in multiple solid tumor tissues, including melanoma and breast cancer (*9–13*). In a study in cervical squamous cell carcinoma, increased TMEM33 expression was associated with poor prognosis and reduced tumor infiltration of dendritic cells as well as CD8^+^ and Th1 CD4^+^ T cells (*9*). Other reported functions of TMEM33 include its role in intracellular Ca^2+^ buffering, identified in proximal convoluted renal tubule cells, where it engages the ER-resident Ca^2+^ channel polycystin-2, promoting ER Ca^2+^ depletion (*14*). Moreover, by supporting the biosynthesis and function of the E3 ligase RNF5 (*15*), TMEM33 contributes to proteostasis and lipid homeostasis via RNF5-mediated ubiquitination and turnover of substrates, such as the sterol-responsive protein, SCAP (*16*).

Given the elevated expression of TMEM33 in multiple cancers and its reported roles in host immunity, we sought to define its function/s within the cancer immunity cycle and how this impacts tumor growth and progression. Using a constitutive *Tmem33^-/-^* mouse model, we found that TMEM33 loss improved tumor control across melanoma models, with enhanced CD8^+^ T cell infiltration and function. Adoptive transfer approaches further revealed that TMEM33 functions intrinsically within CD8^+^ T cells to govern the accumulation of progenitor exhausted TCF-1^+^PD-1^+^ (Tpex) populations upon tumor challenge, a subset widely recognized to sustain long-term CD8^+^ T cell immunity and promote responsiveness to immunotherapy (*17–22*). TMEM33 deletion also enhanced the efficacy of *ex vivo*-expanded tumor-specific CD8^+^ T cells, suggesting broader relevance for adoptive cell therapies. Finally, analysis of a large, prospectively recruited melanoma patient cohort revealed that reduced CD8^+^ T cell-specific TMEM33 expression associated with improved clinical outcomes. Collectively, these findings highlight TMEM33 as a novel immunoregulatory factor in scaling CD8^+^ T cell anti-tumor responses.

## RESULTS

### Deletion of host TMEM33 improves tumor control

To begin to evaluate the impact of host TMEM33 expression on tumor progression, we leveraged a constitutive *Tmem33^-/-^* C57BL/6 mouse model developed previously (*14*). Following subcutaneous challenge with the ovalbumin (‘OVA’)-expressing mouse melanoma cell line, B16F10 (‘B16F10-OVA’), *Tmem33^-/-^* female mice exhibited significantly delayed tumor growth by day 14, and improved survival compared to wildtype (WT) controls (**Fig. 1A; Supplementary Fig. 1A**). Tumor growth was similarly abrogated in male mice (**Supplementary Fig. 1B**). When challenged with YUMM1.7-OVA tumors (**Fig. 1B**), a melanoma model more closely reflecting the mutational landscape of human disease (*Braf^V600E^*, *Pten^−/−^*, *Cdkn2a^−/−^*), *Tmem33^-/-^* mice were afforded similar protection. To ascertain underlying immunological events governing tumor control, we next isolated CD45^+^ fractions from B16F10-OVA tumors harvested at volume endpoint (≤1000 mm^3^) and analyzed these using flow cytometry. In accordance with their restricted tumor growth, tumors from *Tmem33^-/-^* mice had a higher proportion of CD8^+^ T cells compared to WT controls, whilst the CD4^+^ compartment was not significantly different **(Fig. 1C, D**). Greater frequencies of *Tmem33^-/-^*CD8^+^ tumor-infiltrating lymphocytes (TILs) expressed PD-1 relative to WT (**Fig. 1E, F**), potentially signifying an increase in antigen engagement, although PD-1 expression among CD4^+^ cells was comparable between genotypes. Inoculating with MC38 tumors (colorectal adenocarcinoma), we also observed elevated intratumoral CD8^+^ T cell frequencies in *Tmem33^-/-^* mice compared to WT whilst CD4^+^ compartments were unaltered, as was seen for both B16F10-OVA and YUMM1.7-OVA tumors (**Supplementary Fig. 1C**). Notably however, this was not accompanied by enhanced tumor control (**Supplementary Fig. 1D**), potentially reflecting differences in antigenic breadth and weaker dominance of tumor-specific CD8^+^ responses compared with OVA-expressing models.

**Figure 1.**
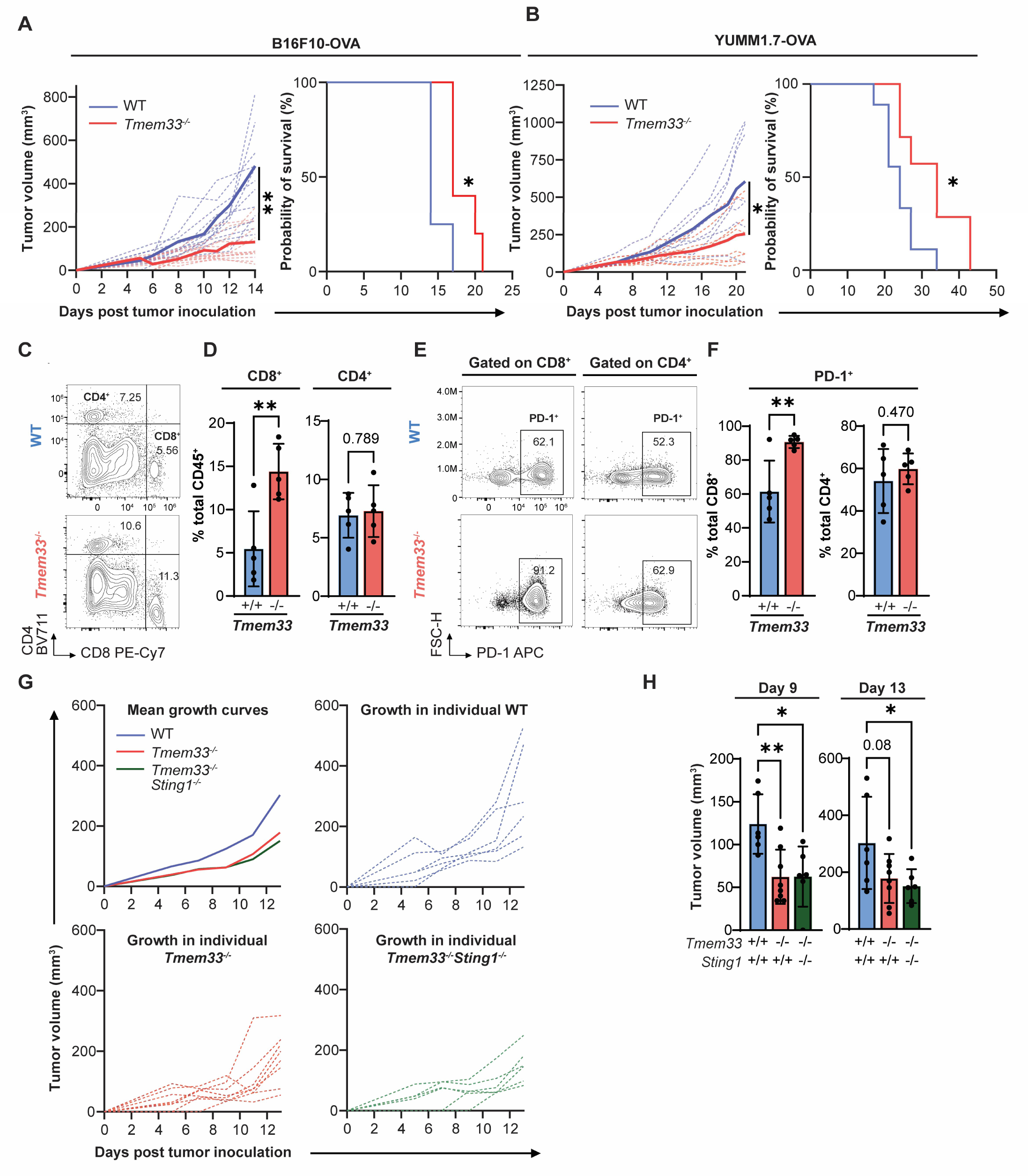
TMEM33 deletion in mice associates with improved tumor control and enhanced CD8^+^ T cell infiltration. **A**, B16F10-OVA tumor growth monitoring in WT and *Tmem33^-/-^*mice at indicated timepoints following cell inoculation (left). Solid lines, mean tumor volumes; dashed lines, tumor volumes for individual mice. Volumes were compared at day 14 post-implantation. Kaplan-Meier survival curves are indicated for equivalent mice (right), compared using the log rank test (Mantel-Cox). N = 8 mice/genotype. **B**, Equivalent analyses as in **A** for YUMM1.7-OVA tumors. Volumes were compared at day 21 post-implantation. N = 8 mice/genotype. **C**, Representative flow cytometry plots indicating CD8^+^ and CD4^+^ populations among total CD45^+^ fractions of B16F10-OVA tumors harvested from WT and *Tmem33^-/-^* mice, with percentage values indicated. **D**, Quantification of CD8^+^ and CD4^+^ frequencies represented in **C**. N = 5 mice/genotype. **E,** Representative plots indicating PD-1^+^ cells among B16F10-OVA tumor CD8^+^ (left) and CD4^+^ (right) compartments, with percentage values indicated. **F**, Quantification of PD-1^+^ cells among total CD8^+^ (left) and CD4^+^ (right) populations. **G**, B16F10-OVA tumor growth in WT, *Tmem33^-/-^* and *Tmem33^-/-^Sting1^-/-^*mice. Mean growth profiles are indicated for each mouse genotype, alongside curves for individual mice. Volumes were compared at indicated timepoints using one-way ANOVA with post-hoc Dunnett’s tests. Unpaired two-tailed student’s *t* tests were performed for tumor volume comparisons in **A** and **B**, and for cell population analyses in **D** and **F**. Bars and error represent mean and SD. \*\**p≤*0.01; \**, p≤*0.05; *p>*0.05 numerically indicated.

Considering the documented roles for TMEM33 in negatively modulating STING (*8*, *23*), and the importance of host STING in triggering anti-tumor immunity (*24*), we asked if the tumor protective effects conferred by *Tmem33* deletion in mice were STING-dependent. To achieve this, we generated *Tmem33^-/-^Sting1^-/-^*mice (**Supplementary Fig. 1E, F**) and assessed B16F10-OVA tumor growth in this novel model (**Fig.1G, H**;). Restricted tumor progression observed in *Tmem33^-/-^* mice was similarly evident in *Tmem33^-/-^Sting1^-/-^* mice, suggesting that the capacity of TMEM33 to scale anti-tumor immune responses operates independently of STING.

To ascertain whether the improvement to CD8^+^ T cell tumor infiltration observed in *Tmem33^-/-^* hosts was attributable to baseline immunological differences, we quantified splenic T cell compartments in tumor-naive mice. The total populations of CD3^+^, CD4^+^ and CD8^+^ T cells were comparable between genotypes (**Supplementary Fig. 2A, B**). Moreover, thymic T cell development, assessed by the abundance of single positive CD4^+^, single positive CD8^+^, and double positive thymocytes, was unaltered (**Supplementary Fig. 2C**). These data suggest that TMEM33 deletion does not perturb basal T cell homeostasis and its immunoregulatory effects are likely restricted to contexts of immune challenge.

### Anti-tumor CD8^+^ T cell functionality is augmented in *Tmem33^-/-^* mice

Tumor-targeting CD8^+^ T cell function crucially underpins effective anti-tumor immunity. To examine the potential role/s of TMEM33 in modulating intratumoral T cell fate, we used bulk RNA-seq to transcriptomically profile CD8^+^ tumor infiltrating lymphocytes (TILs) magnetically isolated from day 14 B16F10-OVA tumors grown in WT and *Tmem33^-/-^* hosts (**Fig. 2A**). Gene set enrichment analysis (GSEA) revealed significant upregulation of biological processes linked to heightened activation in TILs of *Tmem33^-/-^* mice, including cell cycle progression, DNA metabolism and T cell receptor (TCR) signal modulation (**Fig. 2B**). This was accompanied by a reduction in signatures associated with recognized TMEM33 functions, such as lipid and cholesterol metabolism, Ca^2+^ handling, and apoptotic regulation. Hierarchical clustering and differential expression analysis showed co-enrichment of cytotoxic and effector transcripts from TILs of *Tmem33^-/-^*mice (including *Tnf, Gzmb, Gzmc, Gzme, Gzmf, Prf1, Fasl* and *Ifng*), together with increased expression of *Tbx21* encoding the transcription factor and effector regulator T-bet (*25*), as well as markers of terminal differentiation *(Tox, Lag3, Ctla4* and *Pdcd1*) (**Fig. 2C; Supplementary Fig. 3A**). The transcriptomic data therefore suggest that the absence of TMEM33 improves effector activation and tumoricidal activity among TILs.

**Figure 2.**
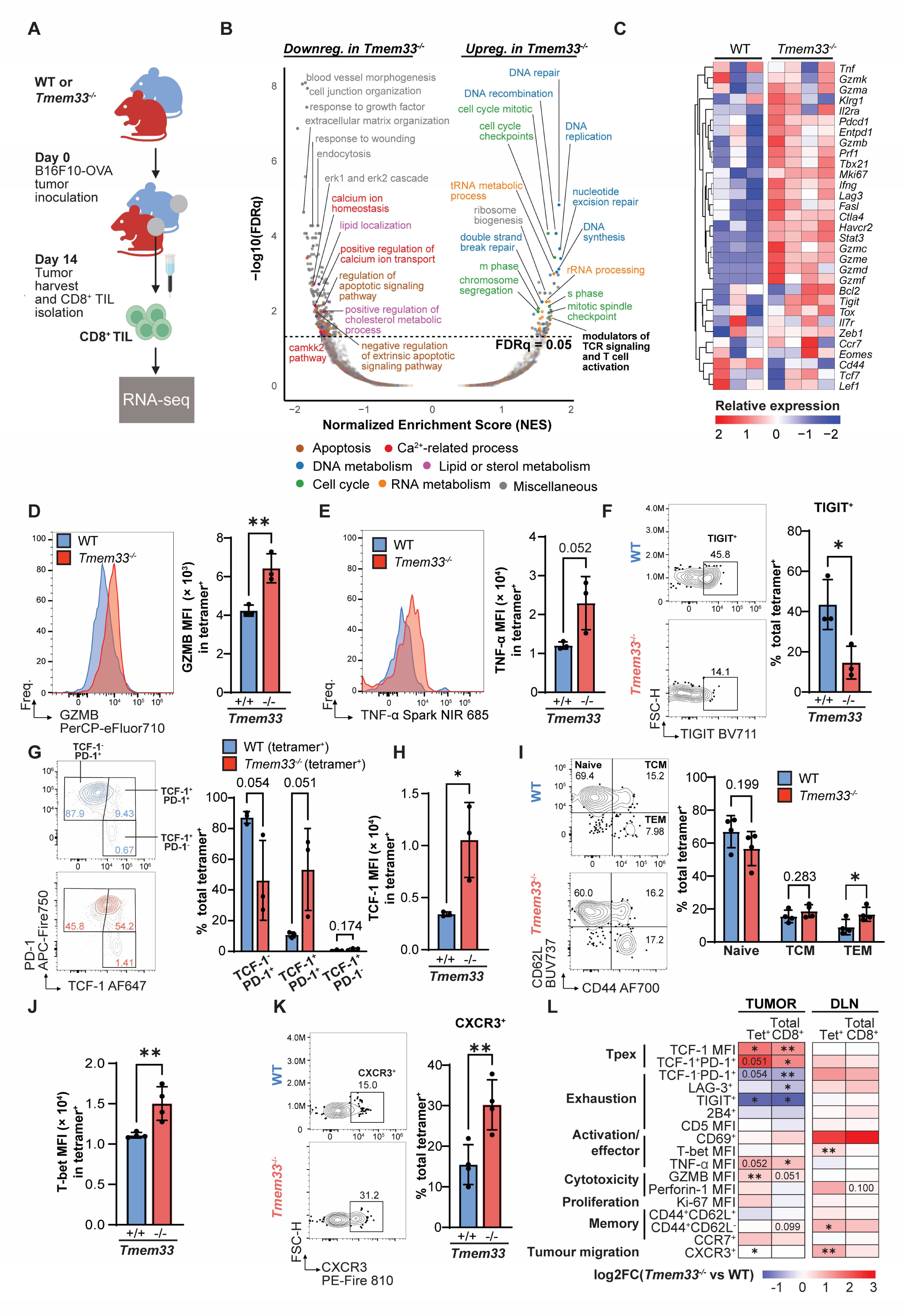
Anti-tumor CD8^+^ T cell fitness is improved in *Tmem33^-/-^* mice. **A**, Schematic indicating experimental workflow and timelines for bulk RNA-seq analysis of CD8^+^ TILs in B16F10-OVA tumors. Designed using BioRender. Tumors were harvested at day 14 following tumor inoculation, and CD8^+^ TILs magnetically labeled and isolated from cell suspensions using MACS columns, prior to bulk RNA-seq. **B**, Volcano plot showing gene set enrichment analysis (GSEA) outputs of *Tmem33^-/-^* CD8^+^ TILs relative to WT counterparts, utilizing GO:BP, CP:BIOCARTA, CP:KEGG, CP:PID, CP:WIKIPATHWAYS, and CP:REACTOME databases. Signatures are plotted according to significance, as −log10 False discovery rate (FDR)-adjusted *p* values (q), vs normalized enrichment score (NES). Dashed line represents significance threshold (FDRq = 0.05). Individual gene sets are colored according to general function **C**, Clustered heatmap of selected genes relating to T cell differentiation, colored according to scaled expression (Z-score). N = 3-4 mice/genotype. **D-K**, Flow cytometric profiling of OVA_257-264_ tetramer^+^ CD8^+^ T cells in day 14 B16F10-OVA tumors or DLNs. For tumor analysis, CD45^+^ immune fractions were magnetically labeled and isolated from harvested tumors prior to staining. Representative contour plots (including labeled percentages of total tetramer^+^) or histograms and adjacent quantification of individual samples are shown for each population or marker of interest. **D**, Granzyme-B (GZMB) and **E,** TNF-α expression in tetramer^+^ TILs (MFI, mean fluorescence intensity). **F**, Proportion of TIGIT^+^ cells among tetramer^+^ TILs. **G**, Tetramer^+^ populations stratified by TCF-1 and PD-1 expression. TCF-1^-^PD-1^+^, exhausted (Tex); TCF-1^+^PD-1^+^, progenitor exhausted (Tpex); TCF-1^+^PD-1^-^, quiescent/memory. In flow plots, tetramer^+^ cells (WT, blue; *Tmem33^-/-^*, red) are superimposed onto respective total CD8^+^ populations (gray). **H,** Mean TCF-1 expression (MFI) in tetramer^+^ TILs. **I**, Characterization of naive (CD44^-^CD62L^+^), central memory (TCM; CD44^+^CD62L^+^) and effector memory (TEM; CD44^+^CD62L^-^) subsets among tetramer^+^ cells in DLNs. **J**, T-bet expression (MFI) among tetramer^+^ cells within DLNs. **K**, Proportion of CXCR3^+^ cells among tetramer^+^ cells within DLNs. **L**, Summary heatmap representation of differential expression analysis or population abundance for specified markers in tumors and DLNs, based on flow cytometry data. Cell coloring represents log2 fold change (FC) in MFI or population abundance in *Tmem33^-/-^* vs WT mice. All flow cytometry data in **D-K** were analyzed using unpaired two-tailed Student’s *t* tests. Bars and error represent mean and SD. **, *p≤*0.01; \**, p≤*0.05; *p>*0.05 values are numerically indicated. For **D-K**, N = 3-4 mice/genotype.

To validate these observations and further dissect the effects of TMEM33 on distinct tumor-specific CD8^+^ T cell compartments, we analyzed day 14 CD45^+^ B16F10-OVA tumor fractions and draining lymph nodes (DLNs) using a high dimensional T cell-focused spectral flow cytometry panel (**Fig. 2D-L; Supplementary Fig. 3B, C**). Consistent with our bulk RNA-seq analysis, granzyme-B and TNF-α were elevated in antigen-specific (OVA_257-264_ tetramer^+^) *Tmem33^-/-^* TILs (**Fig. 2D, E**), indicative of improved cytotoxicity and functionality. This was accompanied by reduced surface expression of inhibitory receptors TIGIT and LAG-3 among both antigen-specific and total CD8^+^ compartments (**Fig. 2F; Supplementary Fig. 3D, E, F**), suggesting delayed exhaustion compared to WT.

CD8^+^ TILs subdivided into three distinct populations based on TCF-1 and PD-1 expression (**Fig. 2G; Supplementary Fig. 3G, H**), with virtually all tumor-specific cells expressing PD-1 owing to antigen exposure. Progenitor exhausted (‘Tpex’) TCF-1^+^PD-1^+^ TILs, which self-renew and proliferatively expand upon ICB treatment to sustain tumor responses (*18–21*, *26*), were enriched in *Tmem33^-/-^* mice, whilst more terminally exhausted TCF-1^-^PD-1^+^ populations were diminished, as observed for antigen-specific (**Fig. 2G**) and total CD8^+^ fractions (**Supplementary Fig. 3H**). In line with Tpex skewing, overall TCF-1 expression was significantly elevated in *Tmem33^-/-^* tetramer^+^ TILs (**Fig. 2H**).

Improvement to *Tmem33^-/-^* CD8^+^ TIL functionality was corroborated using orthogonal unbiased dimensionality reduction analysis of intratumoral T cell compartments (CD3^+^, **Supplementary Fig. 4**). Metaclusters 1-4, representing activated CD8^+^ populations (PD-1^hi^, **Supplementary Fig. 4A-D**), were generally enriched for TCF-1 and granzyme-B in *Tmem33^-/-^* hosts and showed reduced TIGIT expression (**Supplementary Fig. 4E**). In DLNs of the same B16F10-OVA tumor-bearing *Tmem33^-/-^* mice, a greater proportion of antigen-specific CD8^+^ T cells adopted an effector memory phenotype (TEM; **Fig. 2I**) and expressed higher levels of T-bet (**Fig. 2J**). CXCR3 was also more abundant among tetramer^+^ cells (**Fig. 2K**), suggesting a greater responsiveness to tumor-derived chemokine gradients, enabling effective tumor infiltration (*27–30*). In summary, flow cytometric profiling revealed a coordinated shift in tetramer^+^ and total CD8^+^ TIL and DLN populations from *Tmem33^-/-^* mice towards a functionally enhanced, less exhausted phenotype, with improved maintenance of the tumor Tpex compartment (**Fig. 2L**).

### TMEM33 loss does not alter dendritic cell abundance or antigen presenting capacity

To assess whether TMEM33 extrinsically impacts CD8^+^ T cell anti-tumor immunity by influencing dendritic cell (DC)-mediated priming, we compared DC abundance and activation in DLNs from B16F10-OVA tumor-bearing WT and *Tmem33^-/-^* mice at day 14 (**Supplementary Fig. 5A, B**). However, we found that the amount of total DCs, as well as cDC1, cDC2 and migratory subsets did not differ between genotypes (**Supplementary Fig. 5C, D**). At later tumor stages, total and activated tumor DC frequencies were also comparable in *Tmem33^-/-^* mice, although cDC1 abundance was reduced (**Supplementary Fig. 5E, F**), likely reflecting heightened CD8^+^ T-cell activation and subsequent cDC1 clearance (*31*, *32*). To evaluate whether TMEM33 affects antigen cross-presentation, OVA-pulsed bone marrow-derived DCs from WT and *Tmem33^-/-^* mice were co-cultured with the H2-Kb OVA_257-264_ restricted B3Z reporter line (**Supplementary Fig. 5G**) (*33*). TMEM33 deletion did not alter DC cross presentation at either high or low OVA doses (**Supplementary Fig. 5H**), and given the absence of DC differences *in vivo*, we focused future studies on deciphering potential intrinsic effects of TMEM33 within T cell populations.

### *Tmem33^-/-^* naive CD8^+^ T cells respond more potently to polyclonal stimulation

To initially gauge a potential CD8^+^ T cell-intrinsic modulatory role for TMEM33, we analyzed early-activation marker expression and cytokine release in purified naive WT and *Tmem33^-/-^* splenic CD8^+^ T cells stimulated *ex vivo*. The proportions of cells positive for TNF-α and IFN-γ were found to be significantly greater post-stimulation in *Tmem33^-/-^* CD8^+^ T cells compared to WT, whilst baseline frequencies were comparable between strains (**Fig. 3A, B; Supplementary Fig. 6A, B**). Stimulated expression levels of granzyme-B, in addition to surface activation markers CD69 and CD44, were moderately but significantly enhanced among *Tmem33^-/-^* cells (**Fig. 3C-E)**, whilst CD137 and CD25 profiles remained unchanged between genotypes (**Fig. 3F, G**). Equivalent flow cytometric analyses of CD8^+^ compartments among whole splenocytes largely mirrored these findings (**Fig. 3H**). In contrast, parallel assessment of total splenic CD4^+^ T cells revealed more limited differences between WT and *Tmem33^-/-^* samples, with a significant increase only observed for the early activation marker, CD69 (**Fig. 3H**). Together, these initial findings support T cell-specific functions for TMEM33, which are more prominent in the CD8^+^ compartment.

**Figure 3.**
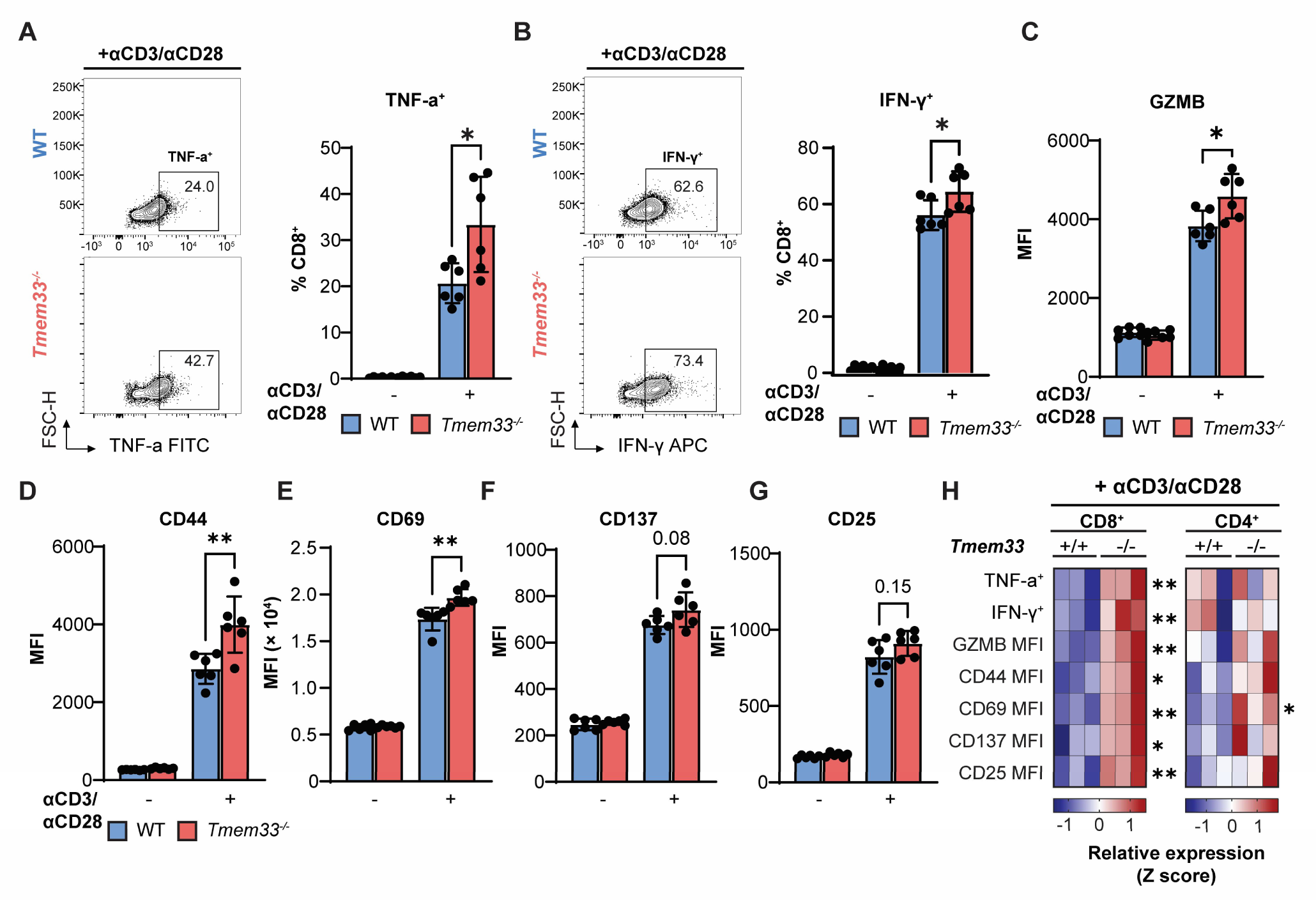
TMEM33 deficiency enhances CD8^+^ T cell responses to polyclonal stimulation. **A-G**, Naive CD8^+^ T cells purified from splenocytes of unchallenged WT and *Tmem33^-/-^* mice were stimulated with anti-CD3/CD28 Dynabeads® and recombinant IL-2 for 24 hours prior to 5 hour incubation with brefeldin-A and flow cytometric analyses of activation markers. **A**, Representative flow plots showing TNF-α expression among stimulated naive CD8^+^ cells (percentages of total CD8^+^ cells indicated), with adjacent quantification of TNF-α^+^ populations among resting and stimulated cells for all mice. **B,** Equivalent analysis (as in **A**) for IFN-γ production. **C,** Expression analysis (mean fluorescent intensity, MFI) for granzyme-B (GZMB), **D**, CD44**, E**, CD69, **F**, CD137 and **G**, CD25. N = 6 mice/genotype. **H**, Heatmap representation of relative expression levels (Z score) for specified markers in CD8^+^ and CD4^+^ populations from WT and *Tmem33^-/-^*whole splenocytes (not purified naive CD8^+^ cells as in **A-G**). Columns indicate individual samples. N = 3 mice/genotype. All data were analyzed using unpaired two-tailed Student’s *t* tests. Bars and error represent mean and SD. **, *p≤*0.01; \**, p≤*0.05; *p>*0.05 values are numerically indicated

### TMEM33 intrinsically modulates Tpex accumulation in tumors and draining lymph nodes, and scales the efficacy of adoptive cell therapy

Given the augmented activity of *Tmem33^-/-^* CD8^+^ T cells observed *ex vivo*, we asked whether TMEM33 intrinsically governs anti-tumor CD8^+^ T cell function *in vivo.* To rigorously test this, we generated allotype-marked (CD45.1^het^) *Tmem33^-/-^*(‘KO’) and *Tmem33^+/+^* (‘WT’) OT-I mice and adoptively transferred their naive CD8^+^ T cells into B16F10-OVA tumor-bearing CD45.2^hom^ hosts (**Fig. 4A**). Tumors and DLNs were harvested on day 13 for flow cytometric analysis. Donor CD45.1^het^ OT-I cells were identified among tetramer^+^ populations co-expressing TCR Vα2 and TCRVβ5 (*34*) (**Supplementary Fig. 7A**). Total TCR Vα2^+^Vβ5^+^ and CD45.1^het^ populations were markedly expanded in tumors of KO OT-I recipients compared to WT counterparts (**Fig. 4B**). These cells exhibited significantly elevated TCF-1 expression, and hence greater Tpex frequency (**Fig. 4C, D**). CTLA-4 expression was significantly reduced among WT OT-I cells (**Fig. 4E**), although the frequencies of cells co-expressing the additional inhibitory receptors LAG-3 and CD39, as well as TIGIT^+^ populations, were equivalent between donor genotypes (**Supplementary Fig. 7B, C**). TNF-α production trended higher in KO OT-I cells, suggesting improved effector function, but did not reach statistical significance (**Supplementary Fig. 7D**). In DLNs, CD45.1^het^ cells from KO OT-I donors were generally more abundant than WT counterparts (**Fig. 4F**; gated from **Supplementary Fig 7E**), and in line with tumor data, showed significantly greater Tpex enrichment (**Fig. 4G**), owing to elevated PD-1 expression (**Fig. 4H**). Transferred cells acquired central and effector memory phenotypes, which were comparable across genotypes (**Supplementary Fig. 7F**). Expression profiles for activation markers CD69 and CD137 were also unchanged (**Supplementary Fig. 7G**).

**Figure 4.**
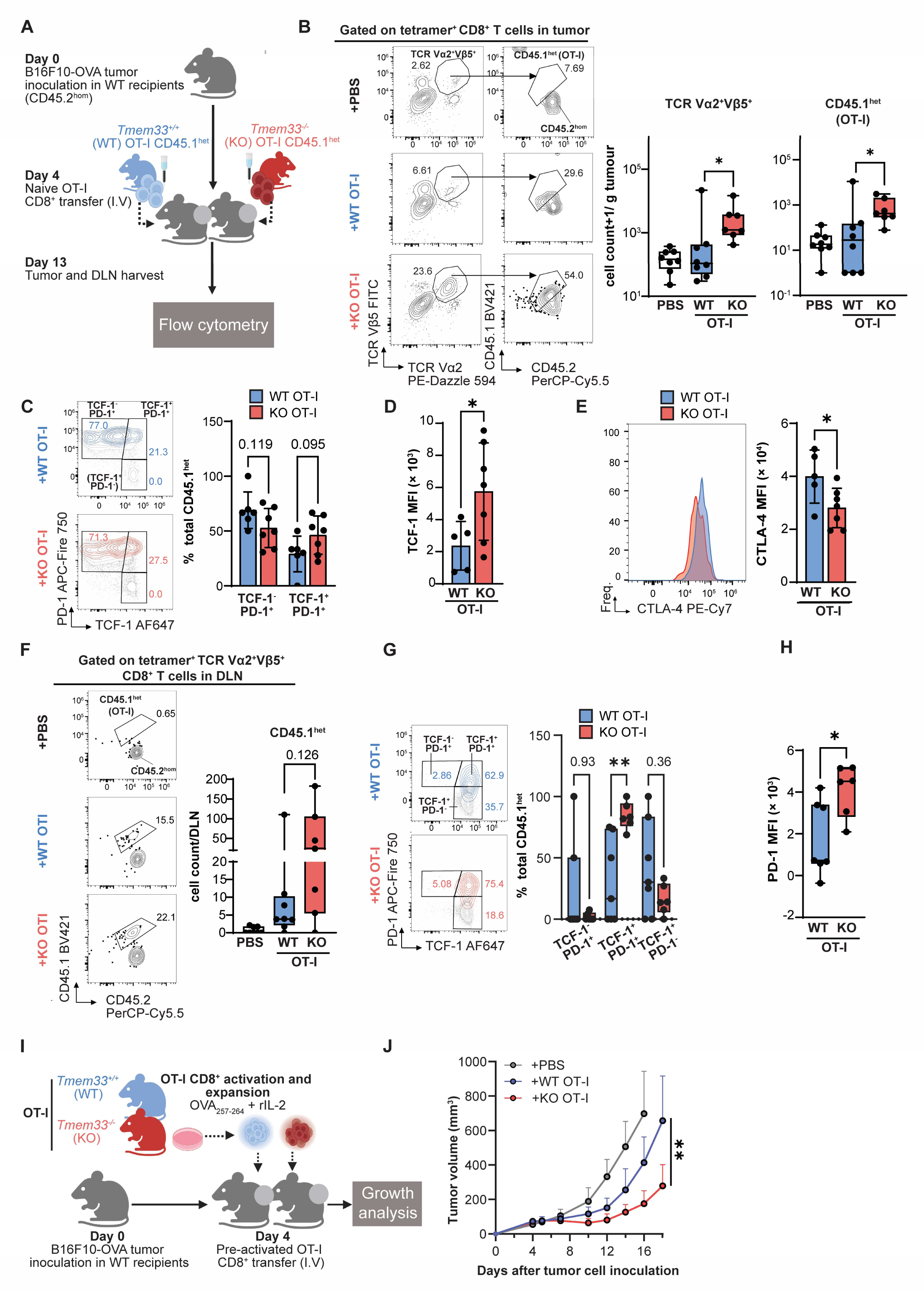
Naive *Tmem33^-/-^*tumor-specific CD8^+^ T cells show enhanced Tpex enrichment and total accumulation upon transfer, while activated *Tmem33^-/-^* CD8^+^ T cells confer superior tumor control. **A**, Schematic outlining adoptive transfer of naïve OT-I CD8^+^ T cells. Wildtype mice (CD45.2^hom^) harboring B16F10-OVA tumors were intravenously administered with naive CD8^+^ cells magnetically isolated (using MACS columns) from *Tmem33^+/+^*or *Tmem33^-/-^* CD45.1^het^ OT-I donors (‘WT’ and ‘KO’, respectively), or PBS (negative control) 4 days post-implantation. Tumors and DLNs were harvested for flow cytometry analysis on day 13. **B**, Representative plots (left) showing TCR Vα^2+^Vβ5^+^ populations among H2-Kb OVA_257-264_ tetramer^+^ CD8^+^ compartments in tumors, and gating on CD45.1^het^ donor cells. Percentages of parent populations are indicated. Absolute OT-I cell counts (per gram of tumor) were determined in tumors of individual recipient mice (right). Cell count+1 is plotted to enable log transformation of 0 values. **C,** Representative plots (left) indicating stratification of intratumoral WT (blue) and KO (red) OT-I cells into subsets according to TCF-1 and PD-1 expression, and their quantification (right). In contour plots, OT-I populations are superimposed onto total CD8^+^ profiles (gray) of recipient mice, and percentage values of total CD45.1^het^ are indicated. **D,** TCF-1 expression among WT and KO OT-I cells in tumors. **E**, CTLA-4 expression among indicated intratumoral OT-I populations with representative histograms for specified markers plotted (left), along with quantified MFI values for individual recipient mice (right). **F,** Representative plots indicating CD45.1^het^ donor cells (as in **B**) among TCR Vα^2+^Vβ5^+^ OVA_257-264_ tetramer^+^ CD8^+^ compartments in DLNs. Absolute cell counts were quantified in each recipient mouse (right). **G,** Representative plots indicating stratification of OT-I cells in DLNs by TCF-1 and PD-1 expression (left; percentage values of total CD45.1^het^ indicated) and quantification of subset frequencies (right) for individual mice. In plots, OT-I populations are superimposed onto respective total CD8^+^ compartments (gray) in recipient mice. **H,** PD-1 expression among WT and KO OT-I cells in DLNs of individual mice. MFI, mean fluorescent intensity. N = 7-8 mice/treatment. Where no absolute CD45.1^het^ cells were identified (cell count = 0), these samples were excluded for downstream functional analysis. **I,** Schematic indicating adoptive transfer of pre-activated OT-I CD8^+^ T cells. Wildtype B16F10-OVA tumor-bearing mice intravenously received *Tmem33^+/+^*(WT) or *Tmem33^-/-^* (KO) OT-I CD8^+^ T cells pre-activated with OVA_257-264_ peptide and recombinant IL-2 (rIL-2), 4 days post-implantation. **J,** Mean growth profiles for B16F10-OVA tumors in mice receiving PBS, WT OT-I or KO OT-I pre-activated CD8^+^ T cells. (left; mean and SD indicated). Volumes were compared at day 18 (unpaired two-tailed Student’s *t* tests). N = 7-8 mice/treatment. For flow cytometry experiments, normality was assessed using Shapiro-Wilk tests. Nonparametric data are presented as box plots (central lines at median, boxes indicating interquartile range and whiskers showing range) and compared using Mann-Whitney *U* tests. For parametric data, bars and error represent mean and SD; data compared using unpaired two-tailed Student’s *t* tests. \**, p≤*0.05; \*\**p≤*0.01; *p>*0.05 numerically indicated. Schematics designed using BioRender.

To determine whether TMEM33 intrinsically shapes CD8^+^ T cell anti-tumor efficacy, we used an adoptive cell therapy approach in which B16F10-OVA tumor-bearing mice received WT or KO OT-I cells following *ex vivo* activation and expansion with OVA_257-264_ peptide and rIL-2 (**Fig 4I**). Strikingly, KO OT-I cells conferred markedly improved tumor control compared with WT OT-I counterparts (**Fig. 4J; Supplementary Fig. 7H, I**). Together, these findings support a T cell-intrinsic role for TMEM33 in governing CD8^+^ T cell fate during tumor challenge, where its deletion favors Tpex enrichment within both tumors and DLNs, and enhances the efficacy of tumor-targeting CD8^+^ T cell therapy.

### TMEM33 deletion increases Ca^2+^ mobilization in naive CD8^+^ T cells but does not affect TCR-induced transients

Elevated intracellular Ca^2+^ is a key downstream feature of TCR stimulation, enabling the transcriptional upregulation of genes essential for T cell activation, proliferation and effector function (*3*). Considering the previously described regulatory functions of TMEM33 in ER Ca^2+^ homeostasis (*14*, *35*) and the Ca^2+^-associated transcriptomic changes observed in intratumoral *Tmem33^-/-^* CD8^+^ T cells (**Fig. 2B**), we hypothesized that TMEM33 may influence CD8^+^ T cell function through changes in intracellular Ca^2+^ dynamics upon activation. Ca^2+^ transients induced by the Ca^2+^ ionophore ionomycin were significantly elevated in *Tmem33^-/-^* naive CD8^+^ T cells (**Supplementary Fig. 8A**). In contrast, TCR-induced Ca^2+^ flux following combined anti-CD3 and anti-CD28 stimulation trended higher in *Tmem33^-/-^* cells but did not reach statistical significance (**Supplementary Fig. 8B**), as was also observed for baseline Ca^2+^ levels (**Supplementary Fig. 8C**). Thus, whilst the altered response to ionomycin aligns with the existing literature in supporting a fundamental role for TMEM33 in moderating intracellular Ca^2+^ buffering, this function alone does not appear to fully account for the enhanced T cell responses measured upon polyclonal stimulation observed previously (**Fig. 3**).

### Reduced CD8^+^ T cell-specific TMEM33 expression associates with improved survival and elevated TCF7 in melanoma patients

To examine the clinical relevance of TMEM33 in governing T cell-mediated tumor protection, we analyzed its expression in peripheral CD8^+^ T cells using bulk RNA-seq data generated from a large, prospectively recruited melanoma patient study cohort described previously (*36*, *37*). CD8^+^ T cells were isolated from the blood of 236 patients with advanced or metastatic melanoma prior to treatment with single agent anti-PD-1 or combination anti-CTLA-4/anti-PD-1 immune checkpoint blockade (ICB; baseline) or following one cycle of ICB. Kaplan-Meier analyses revealed that patients with below-median CD8^+^ T cell-specific *TMEM33* expression at baseline experienced significantly improved overall survival (OS; HR = 1.48, *p* = 0.034, log-rank test; **Fig. 5A**) and progression-free survival (PFS; HR = 1.65, *p =* 0.0021, log-rank test; **Fig 5B**), whereas post-ICB expression only predicted improved OS (HR = 1.55, *p =* 0.042, log-rank test, **Supplementary Fig. 9A**), but not PFS (HR = 1.36, *p =* 0.102, log-rank test, **Supplementary Fig. 9B**). In multivariable Cox models adjusting for age, gender, and *BRAF* status, the baseline association with PFS proved robust (*p* = 0.011, **Fig. 5C**), although OS effects and post-ICB PFS were not statistically significant (**Supplementary Fig. 9C-E**). Consistent with our *in vivo* data showing elevated TCF-1 among intratumoral *Tmem33^-/-^* OT-I cells, the expression of *TCF7* (the human gene encoding TCF-1) in patient CD8^+^ T cells inversely correlated with *TMEM33* transcript abundance both at baseline and following ICB treatment (**Fig. 5D**). Collectively, these data suggest that reduced TMEM33 expression in CD8^+^ T cells predicts clinically favorable outcomes and further support a relationship between TMEM33 and TCF-1-expressing cell populations.

**Figure 5.**
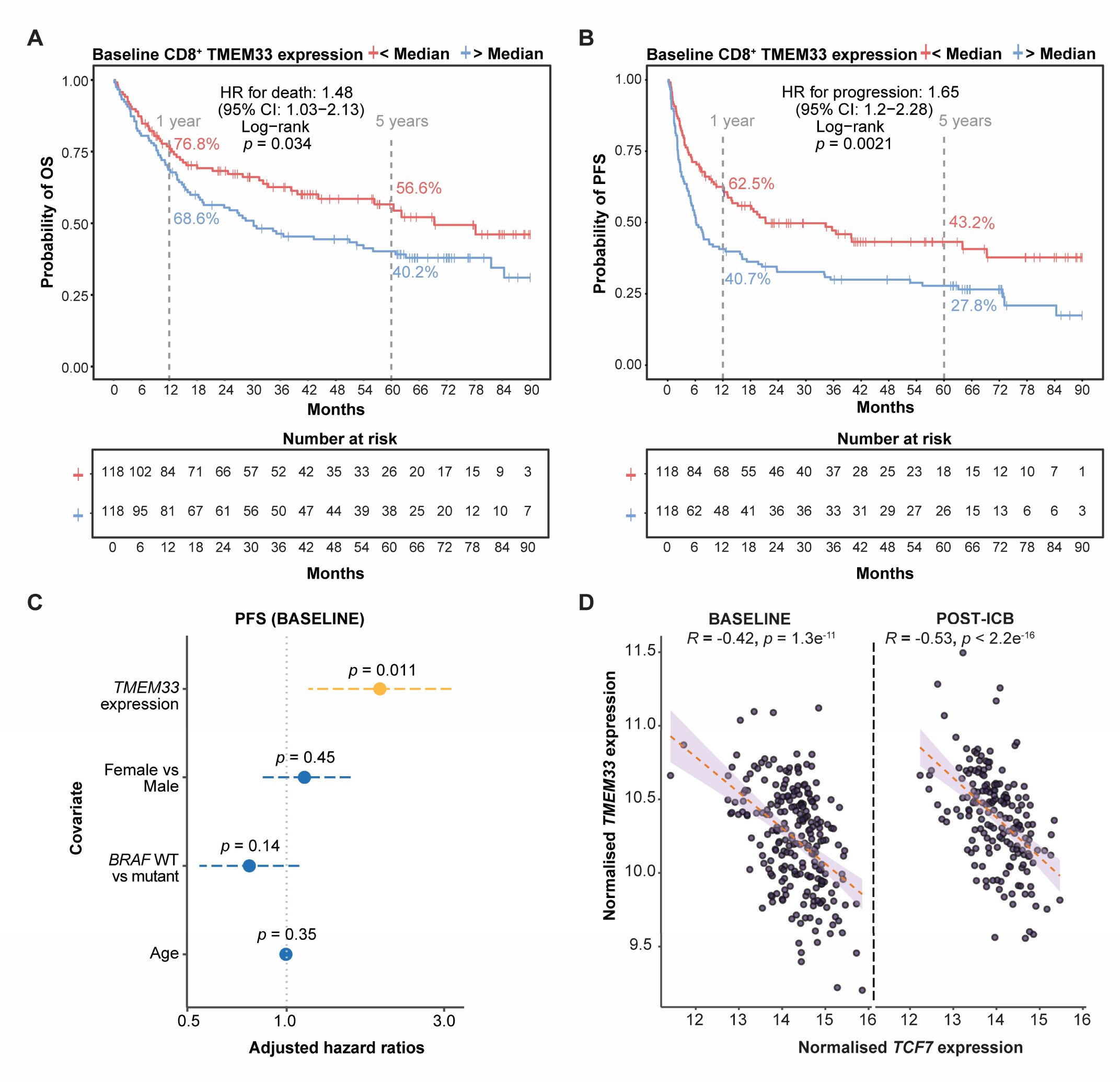
Reduced *TMEM33* expression in melanoma patient CD8^+^ T cells predicts improved survival and correlates with elevated *TCF7* expression. **A**,**B**, Kaplan-Meier analyses indicating overall survival (OS, **A**) and progression-free survival (PFS, **B**) in patients with below- or above-median *TMEM33* expression in peripheral CD8^+^ T cells determined at baseline (prior to immune checkpoint blockade, ICB), by bulk RNA-seq. Two-sided log rank test. N = 118 below-median *TMEM33;* N = 118 above-median *TMEM33*. Hazard ratios (HR) for death (**A**) and progression (**B**) are indicated along with 95% confidence intervals (CI). **C,** Cox proportional hazards model assessing the association between baseline *TMEM33* expression and PFS, adjusting for sex, age and *BRAF* status; Wald test. Dashed lines indicate CI. **D,** Spearman’s rank correlation test of normalized *TMEM33* and *TCF7* expression in patient CD8^+^ T cells at baseline (day 0) or 22 days after receiving one cycle of ICB (‘post-ICB’). Spearman’s *rho* (R) and *p* values derived from a two-sided *t* test are indicated. Red dashed lines show least-squares fit with 95% CI (gray). N = 236.

## DISCUSSION

Advancing current understanding of the fundamental biology underpinning successful anti-tumor CD8^+^ T cell immunity is central to explaining response heterogeneity and guiding the development of new therapeutics. We have identified TMEM33 as a modulator of anti-tumor CD8^+^ T cell responses. Constitutive loss of host TMEM33 improved tumor control and survival in melanoma models, coincident with heightened CD8^+^ TIL abundance, enhanced effector function, reduced exhaustion and enrichment of progenitor-exhausted (Tpex; TCF-1^+^PD-1^+^) populations. Crucially, the observed tumor protection occurred independently of STING, a well-documented mediator of innate anti-tumor responses (*24*). Additionally, the effects of TMEM33 loss uniquely manifested in contexts of immune challenge and did not affect basal T cell homeostasis or development.

We demonstrate that TMEM33 can act intrinsically within CD8^+^ T cells, where its deficiency enhances naive responses to polyclonal stimulation *ex vivo*, and promotes Tpex accumulation in tumors and draining lymph nodes (DLNs) during naive T cell adoptive transfer experiments. Their appearance in both compartments is consistent with reports on both Tpex expansion in DLNs prior to tumor infiltration (*38*), and egress from tumors to DLNs via lymphatic drainage (*39*). Originally defined in contexts of chronic viral infection and later reported in cancer, Tpex harbor self-renewal capacity, yield cytolytic effector progeny and sustain long-term CD8^+^ T cell responses under persistent antigen stimulation (*17*, *18*, *20–22*, *26*), positioning them as plausible mediators of the improved tumor control seen in *Tmem33^-/-^* mice. Moreover, Tpex undergo proliferative bursts in response to immune checkpoint blockade and associate with improved clinical outcomes (*18*, *20*, *26*). Consistently, reduced *TMEM33* expression in melanoma patient CD8^+^ T cells correlated with improved survival and higher *TCF7* levels, suggesting a potential relationship between TMEM33 abundance, TCF-1-centered differentiation, and improved clinical benefit.

An important consideration is the cellular mechanism by which TMEM33 governs Tpex and wider CD8^+^ T cell fate upon tumor antigen exposure, and whether this modulatory role is dominant within the tumor microenvironment or during early priming in DLNs. Given the established roles of TMEM33 in ER Ca^2+^ homeostasis (*14*, *35*) together with the altered Ca^2+^-associated transcriptional profile observed for *Tmem33^-/-^* tumor CD8^+^ T cells, we reasoned that TMEM33 may impact Ca^2+^ -dependent TCR signaling and differentiation. Whilst TCR-driven Ca^2+^ flux in naive CD8^+^ T cells was unaffected by TMEM33 loss *ex vivo* despite enhanced overall activation, these acute stimulation assays are not likely to capture the signaling dynamics and associated TMEM33 functions relevant to chronic tumor antigen exposure. Notably, recent work demonstrated that deficiency of TMEM41B, an ER-embedded Ca^2+^ release channel, maintains T cells in a metabolically activated, stem-like state by promoting ER Ca^2+^ overload, and improves responsiveness to both acute and chronic stimulation (*40*). Whether TMEM33 similarly modulates Ca^2+^ handling to influence T cell fate, despite not functioning as a canonical Ca^2+^ channel itself, remains to be determined.

Additional processes shaped by TMEM33 could potentially contribute to T cell fitness during tumor progression. For example, TMEM33 and its partner E3 ligase, RNF5 act in tandem to fine-tune the cellular response to cholesterol and lipids (*16*), key metabolic drivers of CD8^+^ T cell exhaustion within the immunosuppressive tumor microenvironment (*41*). In line with this, gene sets relating to lipid and sterol homeostasis were differentially enriched in *Tmem33^-/-^* CD8^+^ TILs. However, the tumor-protective and CD8^+^ T cell-potentiating effects documented in *Rnf5^-/-^* hosts have been attributed to non-hematopoietic compartments (*42*), in contrast to the T cell-intrinsic function of TMEM33 we propose here.

Adoptive T cell therapies, including patient-derived TILs and genetically engineered T cells, face challenges in maintaining effector function, resisting exhaustion and persisting within hostile tumor microenvironments (*43–45*). Multiple approaches aim to bolster infused T cell fitness, including pharmacologically targeting glycolytic metabolism (*46*) and chromatin remodeling factors (*47*), as well as genetic deletion of inhibitory receptors and intrinsic regulators of T cell function (*48*, *49*). We show that TMEM33 deletion in *ex vivo-*expanded OT-I T cells enhances their therapeutic fitness in controlling tumor progression upon infusion, further highlighting that TMEM33 intrinsically constrains anti-tumor CD8^+^ responses, and that its intervention may improve the performance and durability of adoptively transferred T cell products in the clinic.

Together, using both murine genetics and patient CD8^+^ transcriptomics, we provide evidence supporting a novel role for TMEM33 in modulating tumor-directed CD8^+^ T cell fate and disease progression. Its intrinsic control of Tpex accumulation is clinically significant, given the importance of this subset in enabling CD8^+^ T cell persistence. Together with our data identifying TMEM33 as a critical regulator of response to adoptive cell therapy, we propose TMEM33 as a potential target to strengthen immunotherapeutic efficacy.

## MATERIALS AND METHODS

### Study design

This study was designed to define the role of TMEM33 in regulating anti-tumor immunity. We used constitutive *Tmem33^-/-^* mice to assess the impact of TMEM33 loss on tumor growth, intratumoral immune composition, and CD8⁺ T cell activation across multiple syngeneic tumor models (B16F10-OVA, YUMM1.7-OVA, and MC38). To interrogate intrinsic effects within CD8⁺ T cells, we conducted adoptive transfer experiments using naive *Tmem33^-/-^* and *Tmem33^+/+^* OT-I CD8⁺ T cells and analyzed their states in tumors and draining lymph nodes. We also monitored tumor growth responses to *Tmem33^-/-^* cell therapy using *ex vivo-*expanded OT-I T cells. These approaches were complemented with *ex vivo* stimulation assays of purified naive CD8⁺ T cells to examine early activation and cytokine responses, and with bulk RNA-seq of tumor-infiltrating CD8⁺ T cells to define transcriptional signatures regulated by TMEM33. Dendritic cell abundance and antigen-presentation capacity were evaluated to test for extrinsic contributions to CD8⁺ T cell priming, and Ca²⁺ flux analyses were performed to assess effects on intracellular Ca²⁺ dynamics associated with TCR signaling. To determine clinical relevance, we analyzed bulk RNA-seq data from peripheral CD8⁺ T cells of patients with advanced melanoma treated with immune checkpoint blockade. Experimental end points for tumor studies were set at defined timepoints or tumor volumes. Tumor-bearing mice were randomly assigned to adoptive transfer groups without investigator blinding. All tumor growth studies were performed with at least 5 biological replicates per experimental group.

### Animal studies

All animal studies were conducted in accordance with the UK Animal (Scientific Procedures) Act of 1986, following approval from the Local Ethics Reviews Committee (University of Oxford). Mice were housed in individually ventilated cages maintained at 22°C, 50% humidity, with 12 hour light/dark cycles, and received additional environmental enrichment. C57BL/6N *Tmem33^-/-^* mice were gifted by Prof. Eric Honoré. These mice harbored the *Tmem33^tm1(KOMP)Mbp^* allele, comprising a LacZ insertion in intron 2-3 of the *Tmem33* gene, deleting coding exons 3-6 (*14*). *Sting1^-/-^* mice provided by Jan Rehwinkel, carried the *Sting1^tm1Camb^* allele, featuring an FRT-flanked neomycin resistance cassette and a loxP site inserted into intron 5, along with a second loxP site introduced into intron 2 (*50*). *Tmem33^-/-^* and *Sting1^-/-^* mice were interbred to generate *Tmem33^-/-^Sting1^-/-^* strains. *Ptprc^a^* (CD45.1) congenic, OT-I TCR^tg^ mice were obtained from Prof. Benoit Van den Eynde (Ludwig Institute for Cancer Research) and interbred with *Tmem33^-/-^* mice to yield *Tmem33^-/-^* or *Tmem33^+/+^*OT-I^het^CD45.1^het^ progeny. All mice were genotyped using Transnetyx.

### Tumor implantation and growth studies

B16F10-OVA-ZsGreen cells (‘B16F10-OVA’; gifted by Prof. Audrey Gérard) and MC38 cells (gifted by Prof. Geoff Higgins) were passaged in DMEM (ThermoFisher, 1965092) supplemented with 10% FBS (ThermoFisher, #A5256701) and 100 U/mL penicillin-streptomycin (ThermoFisher, #15070063). YUMM1.7-OVA cells (gifted by Prof. David Withers) were cultured in DMEM/F-12 (1:1) (1X) + GlutaMAX^TM^-I (ThermoFisher, 31331-028) with added MEM non-essential amino acids (ThermoFisher, #11140050) and penicillin-streptomycin (100 U/ml). All tumor cell lines were maintained in a 37°C incubator, under a humidified atmosphere containing 5% CO_2_, and routinely assessed for mycoplasma contamination. Sex- and age-matched mice (7-12 weeks/old) were anesthetized with 3% gaseous isoflurane and subcutaneously injected in the pre-shaved right flank with 1×10^5^ B16F10-OVA cells, 1×10^5^ MC38 cells, or 4×10^5^ YUMM-OVA cells in a 100 µL 50:50 mixture of phosphate-buffered saline (PBS) and Matrigel (Corning, #CLS354234-1EA). Tumor dimensions were measured using digital calipers, and volumes were determined using the formula 0.5×length×width^2^. Mice were euthanized either at specified time points or volume endpoints (600-1000 mm^3^). Unless otherwise specified, all mice used for tumor studies were female.

### Tissue processing

Resected tumors were minced using a scalpel and digested by incubating in RPMI 1640 supplemented with Collagenase/Hyaluronidase (Stemcell, #17139911) for 40 minutes at 37°C on a shaking platform. For studies requiring intracellular cytokine staining, digestion medium was supplemented with 1X brefeldin-A (BioLegend, #420601). Digested tumors were passed through 70 µm strainers and resuspended in ammonium chloride solution (Stemcell. #07850) for 5 minutes (room temperature, RT) to lyse erythrocytes. Tumor suspensions were washed in PBS, and immune cells isolated using the EasySep™ Mouse CD45 Positive Selection Kit (Stemcell, #18945) according to the manufacturer’s protocol. Spleen, subinguinal lymph nodes, and thymus tissues were pressed through 70 µm strainers before washing in PBS. Splenic samples were hemolyzed using ammonium chloride as previously described. All tissue-derived cell preparations were counted by trypan blue exclusion prior to staining for flow cytometry.

### Naive CD8^+^ T cell isolation and activation

Naive CD8^+^ T cells were magnetically purified from whole splenocytes of unchallenged 7-12 week/old WT and *Tmem33^-/-^* mice (Miltenyi Naive CD8a^+^ T Cell Isolation Kit, mouse, #130-096-543), and transferred to U-bottom 96/well plates (1×10^5^ cells/well) in 200 µL T cell medium: RPMI 1640 supplemented with 10% FBS, MEM non-essential amino acids, 100 U/mL penicillin-streptomycin, 1 mM sodium pyruvate (ThermoFisher, #11360070), 2 mM L-glutamine (ThermoFisher, #25030081) and 50 µM 2-mercaptoethanol (ThermoFisher, #31350010). Cells were administered 7×10^4^ pre-washed anti-CD3/CD28-coupled Dynabeads® (ThermoFisher, #11452D) and 900 U/mL recombinant IL-2 (rIL-2; BioLegend, #575404) for 24 hours (37°C, 5% CO_2_), followed by a 4 hour treatment with brefeldin-A and subsequent staining for flow cytometry. Alternatively, for activation of whole splenocytes, 5×10^6^ cells were seeded per well in T cell medium and treated with 4×10^5^ Dynabeads® and rIL-2 for 24 hours (37°C, 5% CO_2_), prior to brefeldin-A incubation.

### Adoptive transfer experiments

C57BL/6 WT female mice (8 weeks/old; Charles River) were subcutaneously challenged on day 0 with B16F10-OVA tumors as previously described, and randomized according to the order in which they were inoculated. All adoptive T cell transfers were administered in 200 µL PBS via intravenous tail vein injection on day 4 following tumor implantation. For naive T cell transfers, tumor-bearing mice each received 8×10^4^ naive CD8^+^ T cells magnetically isolated from spleens and subinguinal lymph nodes of either *Tmem33^+/+^* or *Tmem33^-/-^* OT-I^het^CD45.1^het^ female mice, as described previously.

For activated T cell transfers (adoptive T cell therapy), splenocytes from OT-I donor mice (*Tmem33^+/+^*or *Tmem33^-/-^*) were first expanded *ex vivo* in T cell medium supplemented with 10 nM OVA_257-264_ peptide and 20 U/mL rIL-2 for 72 hours (37°C, 5% CO_2_), at a starting density of 5×10^5^ cells/mL. Cells were then washed twice in RPMI and cultured at 1×10^6^ cells/mL for a further 24 hours supplemented with IL-2 alone. Expanded CD8^+^ T cells were enriched using Lymphoprep density gradient medium (Stemcell, #18060) as per the manufacturer’s instructions prior to additional washing and resuspension in PBS. A total of 1×10^6^ activated OT-I cells were administered to each tumor-bearing mouse.

### BMDC generation and cross-presentation assay

Bone marrow from femora and tibiae of sacrificed mice were flushed with RPMI 1640 using a syringe and passed through a 70 µm strainer. For the generation of bone marrow-derived dendritic cells (BMDCs), cells were cultured in IMDM media (ThermoFisher, #12440061) supplemented with 10% FBS, 100 U/mL penicillin-streptomycin, 50 µM 2-mercaptoethanol (ThermoFisher, #11528926) 100 ng/mL recombinant FLT3L (ThermoFisher, #250-31L-50UG) and GM-CSF (ThermoFisher, # 315-03-20UG), for 15 days using non-treated culture dishes. BMDCs were transferred to 96-well V bottom plates (5×10^4^ cells/well) and activated with 10 µg/mL poly(I:C) HMW (InvivoGen, #tlrl-pic) for 24 hours. Cells were subsequently treated with 5 µg/mL or 20 µg/mL BioMag®Plus Carboxyl superparamagnetic microparticles (Polysciences, #86010-1) pre-coated with full length chicken ovalbumin (OVA; Invivogen, #vac-stova) or bovine serum albumin for 4 hours. Alternatively, BMDCs were spiked with free-form OVA_257-264_ (SIINFEKL) peptide (Genscript; 100 ng/mL) for the same duration. Antigen-pulsed BMDCs were washed twice with PBS and co-cultured with 5×10^4^ B3Z cells for 24 hours. The B3Z T cell hybridoma line, gifted by Dr Ahmet Hazini, expressed a H2-K_b_:OVA_257-264_-restricted T cell receptor and an NFAT-inducible β-galactosidase construct (*33*). Following incubation, cells were washed twice with PBS. B3Z lysates were prepared and processed using the Promega β-Galactosidase Enzyme Assay System (#E2000), according to the manufacturer’s instructions. Absorbance was measured at 420 nm using a POLARstar OMEGA microplate reader (BMG Labtech).

### Flow cytometry

Following quantification of single cell suspensions, 2×10^6^ cells were transferred to 96-well V bottom plates (Corning) and incubated with TruStain FcX^TM^ (BioLegend, #10139) in PBS (1:50 dilution) for 10 minutes at 4°C to block FcγR binding. Where necessary, cells were stained with PE-conjugated OVA_257-264_ H2-Kb SIINFEKL tetramer (NIH Tetramer Core Facility, Emory University) for 1 hour at 4°C. Cells were then subjected to surface staining either for 30 minutes on ice or 15 minutes at 37°C, followed by incubation with viability dyes in 50 µL PBS (30 minutes, 4°C) or flow cytometry staining buffer (2% FBS, 2 mM EDTA in PBS). All flow cytometry staining reagents and their incubation conditions are specified in **Supplementary Table 1.** Cells were fixed for 30 minutes at 4°C eBioscience^TM^ Foxp3/Transcription Factor Staining Buffer Set (ThermoFisher, #00-5523-00) and washed in Permeabilization buffer. Intracellular staining was performed in Permeabilization buffer at 45 minutes (RT) or overnight (4°C). Cells were washed three times with PBS prior to acquisition on the Cytek™ Aurora 4L (16UV-16V-14B-8R) or LSRFortessa (BD Biosciences), using SpectroFlo or BDFACSDiva (8.0) softwares, respectively. For absolute cell quantification, samples received 5000 counting particles (Spherotech, #ACBP-50-10) immediately prior to acquisition. Flow cytometry data was analyzed using FlowJo v10 or the Spectre package on R v4.4.0 (*51*).

### Western blotting

Harvested splenocytes were rinsed in PBS and solubilized in lysis buffer (150 mM NaCl, 50 mM Tris-HCl pH7.4, 5 mM EDTA) containing 1% Lauryl Maltose Neopentyl Glycol (LMNG, ThermoFisher, #A50940) and cOmplete protease inhibitor cocktail (Roche, #11836170001). Whole cell lysates were clarified by centrifugation (17000×g, 20 min, 4°C) and total protein was quantified by the PierceTM bicinchoninic acid (BCA) assay (ThermoFisher, #23225) according to the manufacturer’s instructions. Protein-adjusted lysates were then heated with 4X Laemmli sample buffer (Biorad, #1610747) supplemented with 10% (v/v) 2-mercaptoethanol (10 minutes, 65°C) and resolved by SDS-PAGE on 4-20% pre-cast polyacrylamide gels (Bio-Rad). Gels were run for 1 hour 25 minutes at 120 V in Tris/Glycine/SDS running buffer using Mini-PROTEAN Tetra electrophoresis cells (Bio-Rad). Proteins were transferred to nitrocellulose membranes using the Trans-Blot Turbo Transfer System (Bio-Rad) and blocked at RT for 1 hour with 5% non-fat dried milk in Tris-buffered saline supplemented with 1% Tween-20 (TBS-T), followed by primary antibody incubation (4°C, overnight) in the same blocking buffer. The following primary antibodies were used in the specified dilutions: anti-TMEM33 (1:1000; Bethyl Laboratories, #A305-597), anti-RNF5 (1:1000; Abcam, #ab154959), anti-STING (1:1000; Cell Signaling Technology, #50494), anti-β-actin (1:5000; GeneTex, #GTX629630). Membranes were washed three times in TBS-T (5 minutes, RT) and incubated with appropriate horse radish peroxidase-conjugated secondary antibodies (1:2000; Horse anti-mouse IgG #7076, and Goat anti-rabbit IgG #7074; Cell Signaling Technology) for 1 hour at RT. After three additional washes in TBS-T, membranes were developed with Immobilon Chemiluminescent HRP substrate of varying strengths (Merck) or SuperSignal^TM^ West Femto Maximum Sensitivity Substrate (ThermoFisher, #34094). Chemiluminescent signals were detected by CCD camera (ChemiDoc MP Imaging System, Bio-Rad).

### Ca^2+^ flux assay

Isolated naïve CD8^+^ T cells were resuspended in T cell medium at a concentration of 1×10^6^ cells/mL. Fluo-4 AM was prepared by combining one part of Fluo-4 AM (ThermoFisher, #F14201) with three parts Pluronic F-127 (ThermoFisher, #P6867) 5% w/v aqueous solution before adding to cells at a final concentration of 2.2 µM and incubating at 37°C for 30 minutes. Cells were then washed twice with Ca^2+^-free DPBS and maintained at 4°C in T cell medium prior to flow cytometric acquisition. Samples were equilibrated at 37°C for 5 minutes before acquiring baseline intracellular Ca^2+^ signals for 50 seconds (BD FortessaTM X-20). Cells were subsequently stimulated for 1 minute with either soluble anti-CD3 (eBioscience, #16-0031085) and anti-CD28 (Tonbo Biosciences, # 70-0281-U500) antibodies, or 10 µg/mL ionomycin (Sigma-Aldrich, #I9657) at 37°C, before re-acquiring for a further 5 minutes. Area under curve (AUC) values were calculated for Ca^2+^ signatures post-stimulation in GraphPad Prism using the trapezoid method.

### Bulk RNA sequencing (RNA-seq) of CD8^+^ tumor-infiltrating lymphocytes (TILs)

Excised tumors were digested and hemolyzed as previously described. Tumor suspensions were magnetically labeled with CD8 (TIL) MicroBeads (Miltenyi, # 130-116-478) before loading onto a MACS Column as described in the manufacturer’s protocol. Isolated CD8^+^ TILs were washed twice in PBS, pelleted and snap-frozen on dry ice. RNA extraction, library preparation, sequencing and initial data processing were conducted by Lexogen NGS Services (Lexogen GmbH, Austria). Total RNA was isolated using the SPLIT RNA Extraction Kit (Lexogen), quantified via UV-Vis spectrophotometry (Nanodrop2000c, ThermoFisher), and integrity verified on a Fragment Analyzer System (Agilent) using the DNF-471 RNA kit (15 nucleotide). Libraries were produced with the QuantSeq 3’ mRNA-Seq Library Prep Kit FWD V2 (Lexogen) and quality assessed using the HS-DNA kit (DNF-474, Agilent). Sequencing was conducted on an Illumina NextSeq2000, generating 100-bp single-end reads, with approximately 5 million reads per sample on average.

Raw reads were evaluated with FastQC for quality metrics, and adapter contamination removed using cutadapt (*52*). Read alignment to the *Mus musculus* reference genome (Ensembl GRCm38.101) was conducted with STAR (*53*). Raw gene counts were determined using featureCounts (*54*), followed by normalization and differential expression using the DESeq2 R package (*55*). Differentially expressed genes were ranked by −Log10(false discovery rate-adjusted *p* value). Gene set enrichment analysis (GSEA) was conducted using the fgsea (v1.28.0) package in R (*56*). Gene sets were obtained via the msigdbr package (v7.5.1) (*57*) and included Hallmark pathways and Canonical Pathways [CP; comprising Kyoto Encyclopedia of Genes and Genomes (KEGG), Reactome, Pathway Interaction Database (PID), WikiPathways and BioCarta databases] from the Molecular Signatures Database (MSigDB) (*58*), in addition to gene sets from the Gene Ontology Biological Process (GO:BP) database (*59*).

### Patient survival and CD8^+^ T cell transcriptomic data analyses

Bulk RNA-seq data for CD8^+^ T cells isolated from peripheral blood of patients in the Oxford Cancer Immunotherapy Toxicity and Efficacy (OxCITE) melanoma cohort (N = 236) were analyzed to assess TMEM33 expression. Cohort recruitment, ethics approval, and detailed methods have been previously described (*36*, *37*). Briefly, patients with advanced or metastatic melanoma eligible for immune-checkpoint blockade were prospectively recruited at Oxford University Hospitals NHS Foundation Trust. Blood was collected and CD8^+^ T cells isolated for bulk RNA-seq analysis at baseline or following one cycle of treatment with either single-agent anti-PD-1 or combination anti-CTLA-4/anti-PD-1. Survival analyses were performed in R (v4.4.0) using the survival (v.3.7-0) and survminer (v.0.4.9) packages. Normalized TMEM33 expression values were determined using DESeq2 and were initially correlated with clinical outcome data (OS and PFS from treatment initiation) via Kaplan-Meier curves with univariate log-rank tests. Multivariable cox proportional hazards models were then conducted adjusting for age, sex and *BRAF* status. Median follow-up time was determined using the reverse Kaplan-Meier method (*60*). Patients with <6 months of follow-up were excluded from the study. Spearman’s rank correlation analysis between *TMEM33* and *TCF7* expression was conducted in R (v4.4.0).

### Statistical analyses

Statistical analyses and graph plotting were performed using GraphPad Prism 9.0 and R (v4.4.0). Data were assessed for normality using Shapiro-Wilk tests, and parametric or nonparametric tests were applied as appropriate. For two-group comparisons, unpaired, two tailed *t* tests (parametric) or Mann-Whitney *U* tests (nonparametric) were employed. Multi-group comparisons were made using ordinary one-way ANOVA (parametric) followed by Dunnet’s post-hoc tests for comparisons to a single control.

Parametric data are graphed as bars representing means with error (standard deviation, SD). Box plots are used for nonparametric datasets, where central lines indicate medians, boxes represent interquartile range, and whiskers highlight maximum and minimum values. *P* values for specified comparisons are indicated on plots as follows: **, *p*≤0.01; *, *p*≤0.05; *p*>0.05 values are numerically indicated. Larger datasets such as RNA-seq were plotted using the R package, ggplot2.

## DECLARATIONS

M.T.J. received doctoral studentship funding from AstraZeneca. E.E.P. has served on advisory boards and performed consultancy for Boehringer Ingelheim, Curadev, InhaTarget, and AkamisBio. She is an employee of the University of Oxford, which has received funding or other research support from AstraZeneca and STIpe Therapeutics. V.P.A. is employed part-time as an Immunology Scientist at Infinitopes. T.E. acts as a consultant for Dark Blue Therapeutics. E.A. has received funding or other research support from Moderna and GSK. A.G. and D.I.G. are employees and shareholders of AstraZeneca. In the last three years, B.P.F. has performed consultancy for NICE Consultancy, Roche, Pathios, UCB, and TCypher, and has received speaker fees from GSK, UCB, and BMS, all outside the submitted work. All other authors declare no competing interests.

## ACKNOWLEDGEMENTS

We thank members of the Parkes, Christianson and Withers laboratories for their practical and conceptual support, and acknowledge Prof. Audrey Gérard for B16F10-OVA-ZsGreen cells, Prof. Geoff Higgins for MC38 cells, Prof. Benoit Van den Eynde for OT-I and CD45.1 congenic strains, and Dr. Ahmet Hazini for the B3Z hybridoma line. We also thank the NIH Tetramer Core Facility (NIH Contract 75N93020D00005; RRID:SCR_026557) for providing the PE-conjugated H2-Kb SIINFEKL (OVA_257-264_) tetramer (#73710), and the Biomedical Services at the University of Oxford for their technical assistance. This work was supported by a Wellcome Clinical Career Development Fellowship to E.E.P. (224623/Z/21/Z); an AstraZeneca-supported studentship to M.T.J.; the John Fell Oxford University Press Research Fund (to E.E.P.); and an MSD Pump Prime Award (to M.T.J.). Support from Cancer Research UK included a studentship to B.B. through the CRUK Oxford Centre (C2195/A31281) and a Senior Cancer Research Fellowship to J.C.C. (C68569/A29217). Prostate Cancer UK provided a Major Award in Immunology supporting J.T.W.K. I.P.P. was supported by the Chinese Academy of Medical Sciences Innovation Fund for Medical Science (2024-I2M-2-001-1); H.E. by NIH grants P01CA275738 and P01CA257804; V.P.A. by the Ludwig Institute for Cancer Research; E.A. by the NIHR Oxford Biomedical Research Centre (NIHR203311); J.R. by the UK Medical Research Council (MR/Y013212/1); and MJWS by an MRC Career Development Award (MR/X020746/1). B.P.F. is funded by a Wellcome Career Development Award (226535/Z/22/Z) and was previously supported by a Wellcome Intermediate Clinical Fellowship (201488/Z/16/Z). Research in the Withers laboratory is funded by a Cancer Research UK Discovery Research Program (DRCNPG-Nov23/100008) and a Wellcome Discovery Award (227890/Z/23/Z). We gratefully acknowledge all patients who participated in this study, and the clinical teams of the Oxford Cancer Centre, the Day Treatment Unit, the Brodey Centre, the Oxford Radcliffe Biobank, and the Churchill Hospital Sample Handling Lab for their invaluable support.

## SUPPLEMENTARY MATERIAL

**Supplementary Table 1.** Reagents used for flow cytometry experiments. #, Number.

| Marker | Fluorochrome | Clone | Supplier | Catalogue # | Working dilution (1/) | Incubation conditions |
| --- | --- | --- | --- | --- | --- | --- |
| B220 | BV650 | RA3-6B2 | BioLegend | 103241 | 100 | 30 mins, 4°C |
| CCR7 | BUV661 | 4B12 | BD Biosciences | 741677 | 50 | 15 mins, 37°C |
| CCR7 | PE-Cy7 | 4B12 | eBioscience | 12-1971-82 | 100 | 1 hr, 37°C |
| CD11b | PE-594 | M1/70 | BioLegend | 101255 | 300 | 30 mins, 4°C |
| CD11c | AF700 | N418 | ThermoFisher | 56-0114-82 | 300 | 30 mins, 4°C |
| CD137 | PE-Cy5 | 17B5 | BioLegend | 106123 | 100 | 15 mins, 37°C |
| CD152 (CTLA4) | PE-Cy7 | UC10-4B9 | BioLegend | 106313 | 170 | 45 mins, RT |
| CD183 (CXCR3) | PE-Fire 810 | S18001A | BioLegend | 155923 | 80 | 45 mins, RT |
| CD244 | BUV563 | C9.1 | BD Biosciences | 749156 | 200 | 15 mins, 37°C |
| CD25 | PE-Dazzle 594 | 3C7 | BioLegend | 101920 | 100 | 30 mins, 4°C |
| CD3 | Spark Blue 550 | 17A2 | BioLegend | 100259 | 200 | 15 mins, 37°C |
| CD3 | APC-Cy7 | 17A2 | BioLegend | 100221 | 100 | 30 mins, 4°C |
| CD3 | BV421 | 17A2 | BioLegend | 100227 | 100 | 30 mins, 4°C |
| CD3 | BUV737 | UCHT1 | BD Biosciences | 612750 | 100 | 30 mins, 4°C |
| CD38 | APC-Fire 810 | 90 | BioLegend | 102745 | 1300 | 15 mins, 37°C |
| CD39 | PE-Fire640 | DuHa59 | BioLegend | 143817 | 200 | 15 mins, 37°C |
| CD4 | BV711 | RM4-5 | BioLegend | 100557 | 100 | 30 mins, 4°C |
| CD4 | BV750 | GK1.5 | BioLegend | 100467 | 1300 | 15 mins, 37°C |
| CD44 | APC-R700 | IM7 | BioLegend | 103025 | 1000 | 15 mins, 37°C |
| CD44 | PE | IM7 | BioLegend | 103007 | 100 | 30 mins, 4°C |
| CD45 | Alexa Fluor 700 | I3/2.3 | BioLegend | 147715 | 100 | 30 mins, 4°C |
| CD45 | PE-Dazzle 594 | 30-F11 | BioLegend | 103145 | 100 | 30 mins, 4°C |
| CD45 | Super Bright 436 | 30-F11 | eBioscience | 62045180 | 170 | 15 mins, 37°C |
| CD45 | BV785 | 30-F11 | BioLegend | 103149 | 400 | 30 mins, 4°C |
| CD45.1 | BV421 | A20 | BioLegend | 110731 | 100 | 15 mins, 37°C |
| CD45.2 | PerCP-Cy5.5 | 104 | BioLegend | 109827 | 200 | 15 mins, 37°C |
| CD5 | FITC | 53-7.3 | BioLegend | 100605 | 25 | 15 mins, 37°C |
| CD62L | BUV737 | MEL-14 | BD Biosciences | 612833 | 640 | 15 mins, 37°C |
| CD69 | BV421 | H1.2F3 | BioLegend | 104527 | 100 | 15 mins, 37°C |
| CD69 | BV605 | H1.2F3 | BioLegend | 147715 | 100 | 30 mins, 4°C |
| CD8 | BUV395 | 53-6.7 | BD Biosciences | 565968 | 100 | 15 mins, 37°C |
| CD8a | PE-Cy7 | 53-6.7 | BioLegend | 100722 | 100 | 30 mins, 4°C |
| CXC3R1 | BV570 | SA011F11 | BioLegend | 149061 | 50 | 15 mins, 37°C |
| EOMES | eFluor450 | Dan11mag | Invitrogen | 48487580 | 50 | 45 mins, RT |
| F4/80 | BV605 | BM8 | BioLegend | 123133 | 100 | 30 mins, 4°C |
| GZMB | PE | QA16A02 | BioLegend | 372208 | 100 | 30 mins, 4°C |
| GZMB | PerCP-eFluor710 | NGZB | Invitrogen | 46889880 | 200 | Overnight, 4°C |
| I-A/I-E (MHC II) | BV510 | M5/114.15.2 | BioLegend | 107635 | 500 | 30 mins, 4°C |
| ICOS | PE-Cy5 | C398.4A | eBioscience | 15-9949-81 | 160 | 15 mins, 37°C |
| IFN-γ | BV480 | XMG1.2 | BD Biosciences | 566151 | 100 | 45 mins, RT |
| IFN-γ | APC | XMG1.2 | BioLegend | 505809 | 100 | Overnight, 4°C |
| IL-2 | BV605 | JES6-5H4 | BioLegend | 503829 | 100 | 45 mins, RT |
| Ki-67 | BV650 | B56 | BD Biosciences | 563757 | 25 | 15 mins, 37°C |
| LAG-3 | BV785 | C9B7W | BioLegend | 125219 | 60 | 15 mins, 37°C |
| Ly108 | BUV496 | 13G3 | BD Biosciences | 750046 | 200 | 15 mins, 37°C |
| Ly6C | BV421 | HK1.4 | BioLegend | 128031 | 500 | 30 mins, 4°C |
| Ly6G | BV650 | 1A8 | BioLegend | 127641 | 400 | 30 mins, 4°C |
| MHC-I (H-2Kb) | PECy5.5 | AF6-88.5 | BioLegend | 116515 | 100 | 30 mins, 4°C |
| NK1.1 | BV650 | PK136 | BioLegend | 108735 | 50 | 30 mins, 4°C |
| PD-1 | APC | 29F.1A12 | BioLegend | 135209 | 100 | 30 mins, 4°C |
| PD-1 | APC-Fire 750 | 29F.1A12 | BioLegend | 135239 | 200 | 15 mins, 37°C |
| PD-L2 | BUV395 | TY25 | BD Biosciences | 752604 | 150 | 30 mins, 4°C |
| Perforin1 | PE-Dazzle 594 | S16009A | BioLegend | 154315 | 25 | 45 mins, RT |
| T-bet | PercCP-Cy5.5 | 4B10 | Invitrogen | 45582580 | 50 | 45 mins, RT |
| TCF-1 | AF647 | C63D9 | CST | 6709S | 1000 | Overnight, 4°C |
| TCR Va2 | PE-Dazzle 594 | B20.1 | BioLegend | 127827 | 200 | 15 mins, 37°C |
| TCR Vβ5 | FITC | MR9-4 | BioLegend | 139513 | 400 | 15 mins, 37°C |
| TIGIT | BV711 | 1G9 | BD Biosciences | 744214 | 500 | 45 mins, RT |
| TNF-α | Spark NIR 685 | MP6XT22 | BioLegend | 506359 | 50 | 45 mins, RT |
| TNF-α | FITC | MP6-XT22 | BioLegend | 506303 | 200 | Overnight, 4°C |
| Viability | Live/Dead Blue | N/A | ThermoFisher | L23105 | 1000 | 30 mins, 4°C |
| Viability | Zombie NIR | N/A | BioLegend | 423105 | 5000 | 10 mins, RT |
| Viability | eFluor780 | N/A | ThermoFisher | 65-0865-14 | 4000 | 30 mins, 4°C |
| XCR1 | AF647 | ZET | BioLegend | 148213 | 150 | 1 hr, 37°C |

**Supplementary Figure 1.**
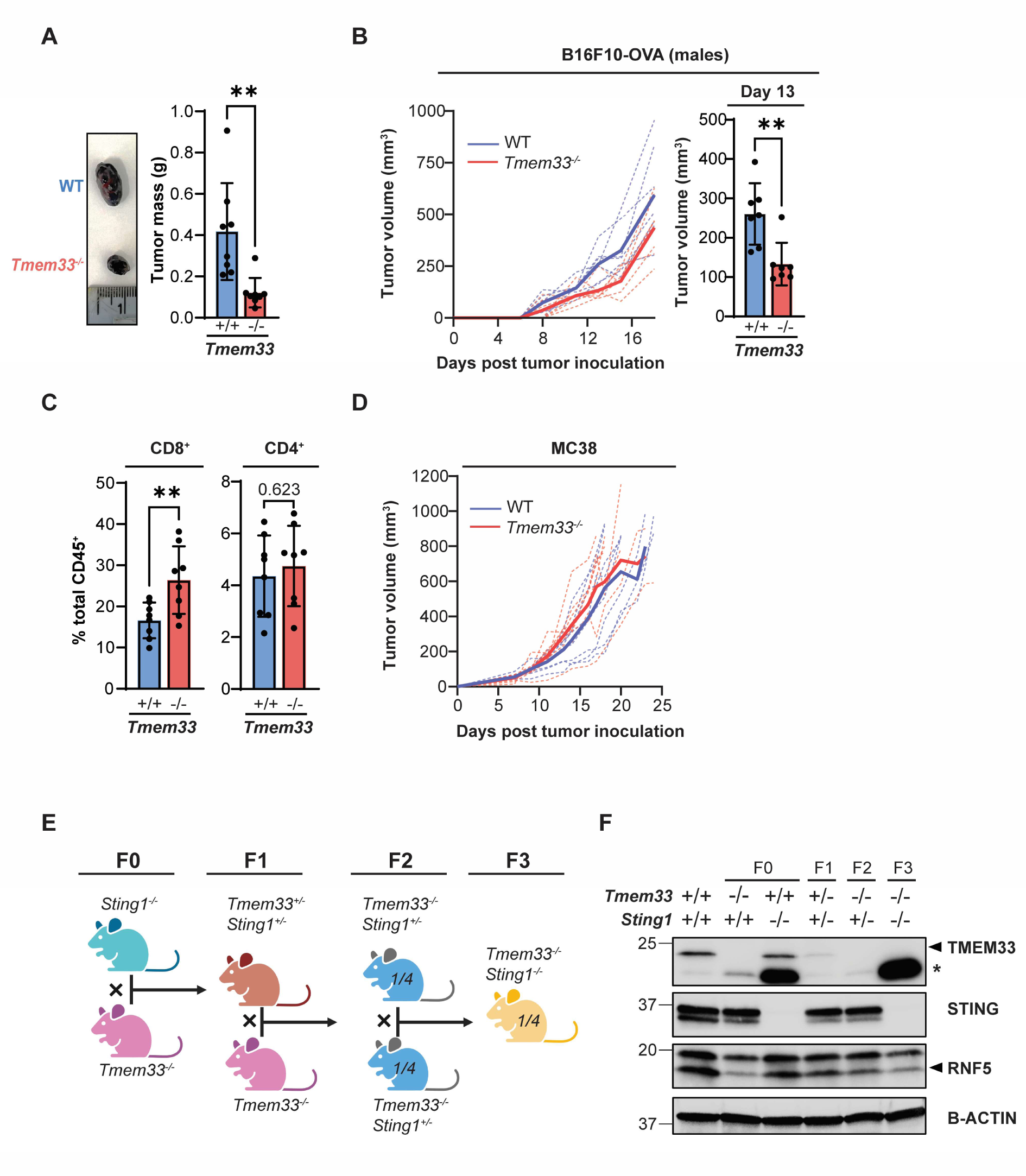
Supporting analyses for Figure 1: TMEM33 deletion in mice associates with improved tumor control and enhanced CD8^+^ T cell infiltration. **A**, Representative photographs (left) and recorded masses (right) of B16F10-OVA tumors resected from WT and *Tmem33^-/-^* mice at day 14 following implantation. N = 8 mice/genotype. **B,** B16F10-OVA tumor growth monitoring in male WT and *Tmem33^-/-^* mice at indicated timepoints following cell inoculation (left). Solid lines, mean tumor volumes; dashed lines, tumor volumes for individual mice. Volumes were compared at day 13 post-implantation. N = 7 mice/genotype**. C**, Quantification of CD8^+^ and CD4^+^ T cell populations within MC38 tumors harvested from WT and *Tmem33^-/-^*mice. Tumors were harvested prior to reaching 1000 mm^3^, and CD45^+^ immune fractions isolated from tumor cell suspensions were analyzed by flow cytometry. N = 8 mice/genotype. **D**, MC38 tumor growth monitoring in WT and *Tmem33^-/-^* mice at indicated timepoints following cell inoculation. Solid lines, mean tumor volumes; dashed lines, tumor volumes for individual mice. **E**, Schematic depicting breeding strategy to generate *Tmem33^-/-^Sting1^-/-^*animals. *Tmem33^-/-^*and *Sting1^-/-^*F0 mice were crossed to generate *Tmem33^+/-^Sting1^+/-^*heterozygotes (F1), which were subsequently mated with *Tmem33^-/-^* F0 mice to yield *Tmem33^-/-^Sting1^+/-^*(F2). F2 mice were interbred to generate *Tmem33^-/-^Sting1^-/-^* double homozygotes. Probabilities of achieving mice harboring a specific genotype within litters are indicated as fractions where relevant. Designed using BioRender. **F,** Western blot analysis of TMEM33, STING and RNF5 expression in F0, F1, F2 and F3 generations of *Tmem33^-/-^*and *Sting1^-/-^*breeding, validating the derivation of *Tmem33^-/-^Sting1^-/-^* mice. *Tmem33* deletion expectedly associated with reduced abundance of its dependent binding partner, RNF5. B-actin was utilized as a loading control. *, non-specific band. For **A-C,** data were compared using unpaired two-tailed Student’s *t* tests; **, *p*≤0.01; *p*>0.05 values are numerically indicated.

**Supplementary Figure 2.**
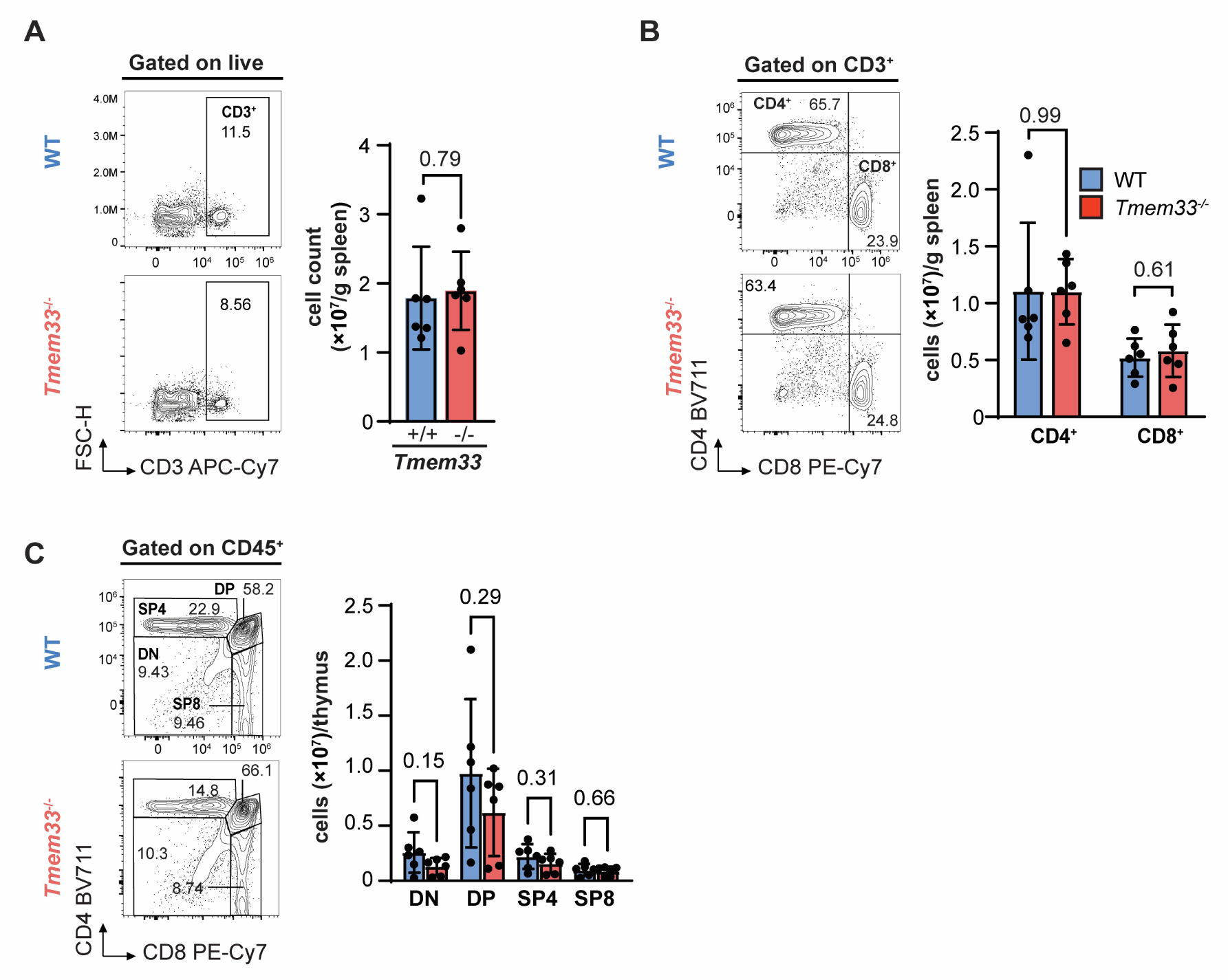
Characterization of splenic T cell and thymocyte populations in unchallenged WT and *Tmem33^-/-^* mice. **A**, Representative flow cytometry plots indicating splenic CD3⁺ T cell populations (percentages of live cells indicated) with adjacent quantification of absolute cell numbers. **B**, Equivalent analyses for splenic CD8⁺ and CD4⁺ T cell subsets (percentages of parent CD3^+^ populations indicated on flow plots). **C**, Representative plots of CD45⁺ thymocytes showing double-negative (DN; CD4⁻CD8⁻), double-positive (DP; CD4⁺CD8⁺), single-positive CD8⁺ (SP8) and single-positive CD4⁺ (SP4) populations (percentages of parent CD45^+^ populations indicated) with adjacent absolute quantification. N = 6 mice/genotype. Mice were 6 weeks at the time of tissue harvest. All data were analyzed using unpaired two-tailed Student’s *t* tests. Bars and error represent mean and SD. *, *p*≤0.05; *p*>0.05 values are numerically indicated.

**Supplementary Figure 3.**
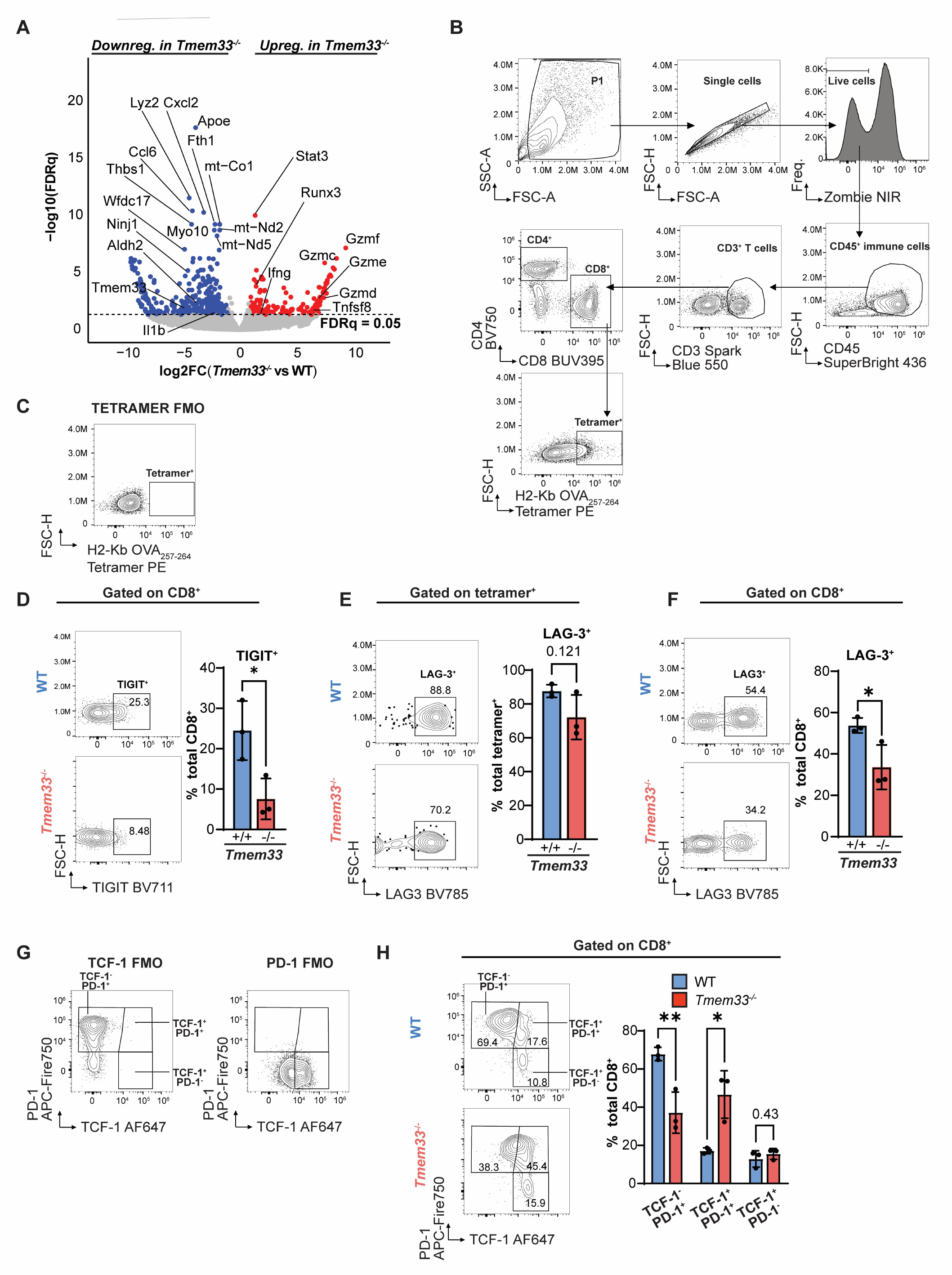
Functional profiling of tumor-specific CD8^+^ T cells, and analysis of total CD8^+^ T cell populations. **A**, Differential gene expression analysis in *Tmem33^-/-^*vs WT CD8^+^ TILs isolated from day 14 B16F10-OVA tumors. Genes are plotted according to significance, as −log10 False discovery rate (FDR)-adjusted *p* values (q), vs log2 fold change (FC) of expression in *Tmem33^-/-^*vs WT CD8^+^ TILs. Red and blue represent significantly upregulated or downregulated genes in *Tmem33^-/-^*TILs, respectively. Gray values are non-significant. Dashed line represents significance threshold (FDRq = 0.05). N = 3-4 mice/genotype. **B**, Full gating strategy employed to identify tumor antigen-specific CD8^+^ T cells, defined by positive staining for the H2-Kb OVA_257-264_ tetramer. **C,** fluorescence-minus-one (FMO) used to define tetramer^+^ cell populations. **D,** Proportion of TIGIT^+^ cells among total tumor CD8^+^ T cells. **E,** Proportion of LAG-3^+^ cells among tetramer^+^ and **F**, total CD8^+^ populations. **G**, FMO controls for TCF-1 and PD-1. **H**, Total tumor CD8^+^ populations stratified by TCF-1 and PD-1 expression. TCF-1^-^PD-1^+^, exhausted (Tex); TCF-1^+^PD-1^+^, progenitor exhausted (Tpex); TCF-1^+^PD-1^-^, quiescent/memory. N = 3 mice/genotype. Data in **D, E, F** and **H** were compared using unpaired two-tailed Student’s *t* tests. Bars and error represent mean and SD. **, *p≤*0.01; \**, p≤*0.05; *p>*0.05 values are numerically indicated.

**Supplementary Figure 4.**
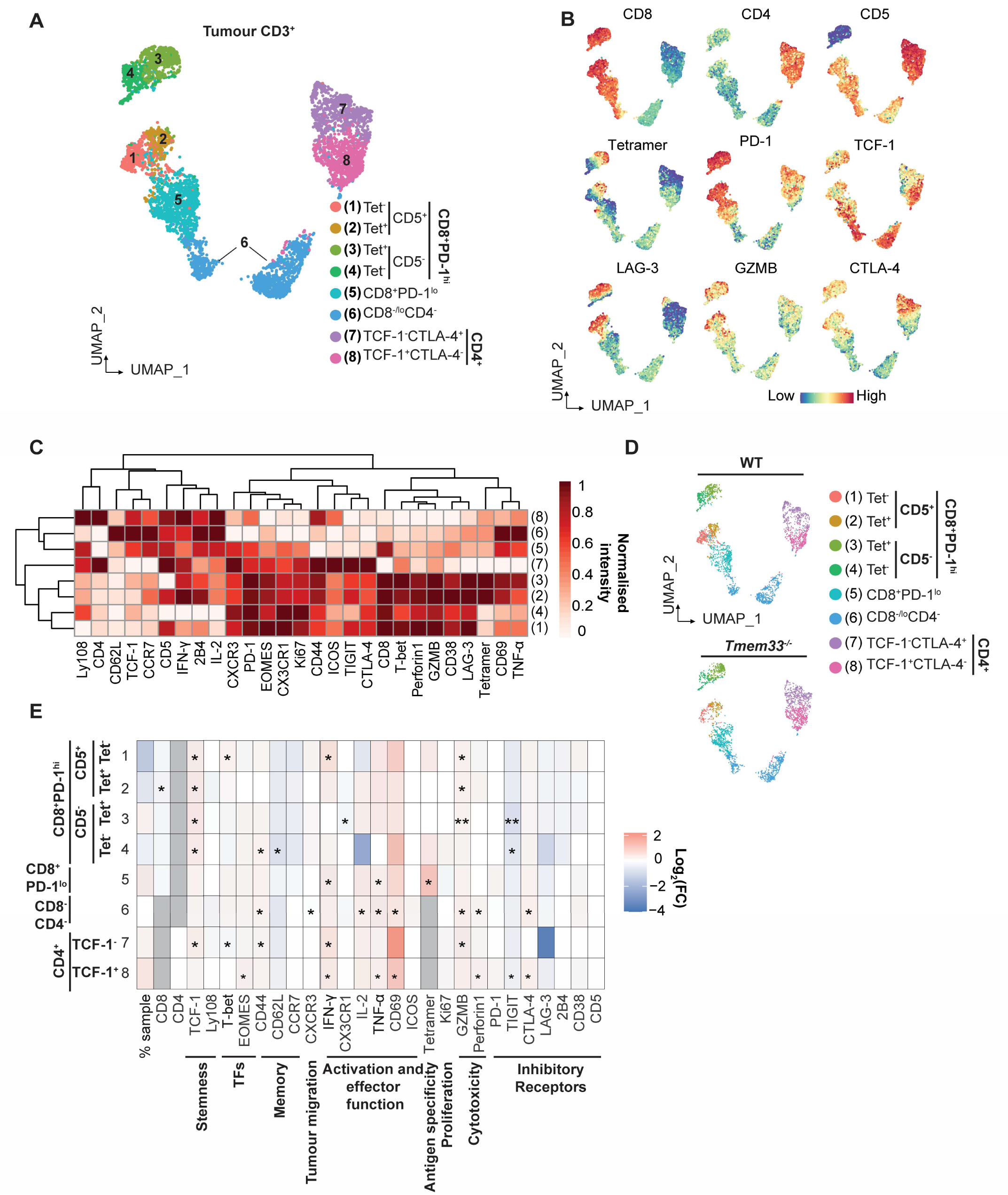
Unbiased dimensionality reduction analysis of intratumoral (B16F10-OVA) T cell populations supports heightened T cell fitness in *Tmem33^-/-^* hosts. UMAP analysis and consensus clustering was performed using Spectre (R Studio) on gated CD3^+^ T cell populations in B16F10-OVA tumors harvested from WT and *Tmem33^-/-^* hosts 14 days post-implantation. **A,** UMAP projection of 6078 CD3^+^ cells concatenated from WT and *Tmem33^-/-^* intratumoral samples. Cells are colored according to their respective metaclusters**. B,** UMAPs highlighting the relative intensity distribution of selected markers across metaclusters. **C,** Consensus clustering of metaclusters according to scaled marker expression. **D,** Visual comparison of metacluster size, displayed as UMAP projections. **E**, Heatmap indicating differential marker expression between mouse strains within metaclusters. The abundance of each metacluster displayed in **D** is highlighted by ‘% sample’. Cell coloring represents log2 fold change (FC) in marker signal in *Tmem33^-/-^* hosts, compared to WT. Gray highlights functionally irrelevant parameters (e.g. CD8 intensity within CD4^+^ metaclusters). TFs, transcription factors. Tet, tetramer. N = 3 mice/genotype. Unpaired two-tailed Student’s *t* tests, \*\**p*≤0.01, \**p*≤0.05.

**Supplementary Figure 5.**
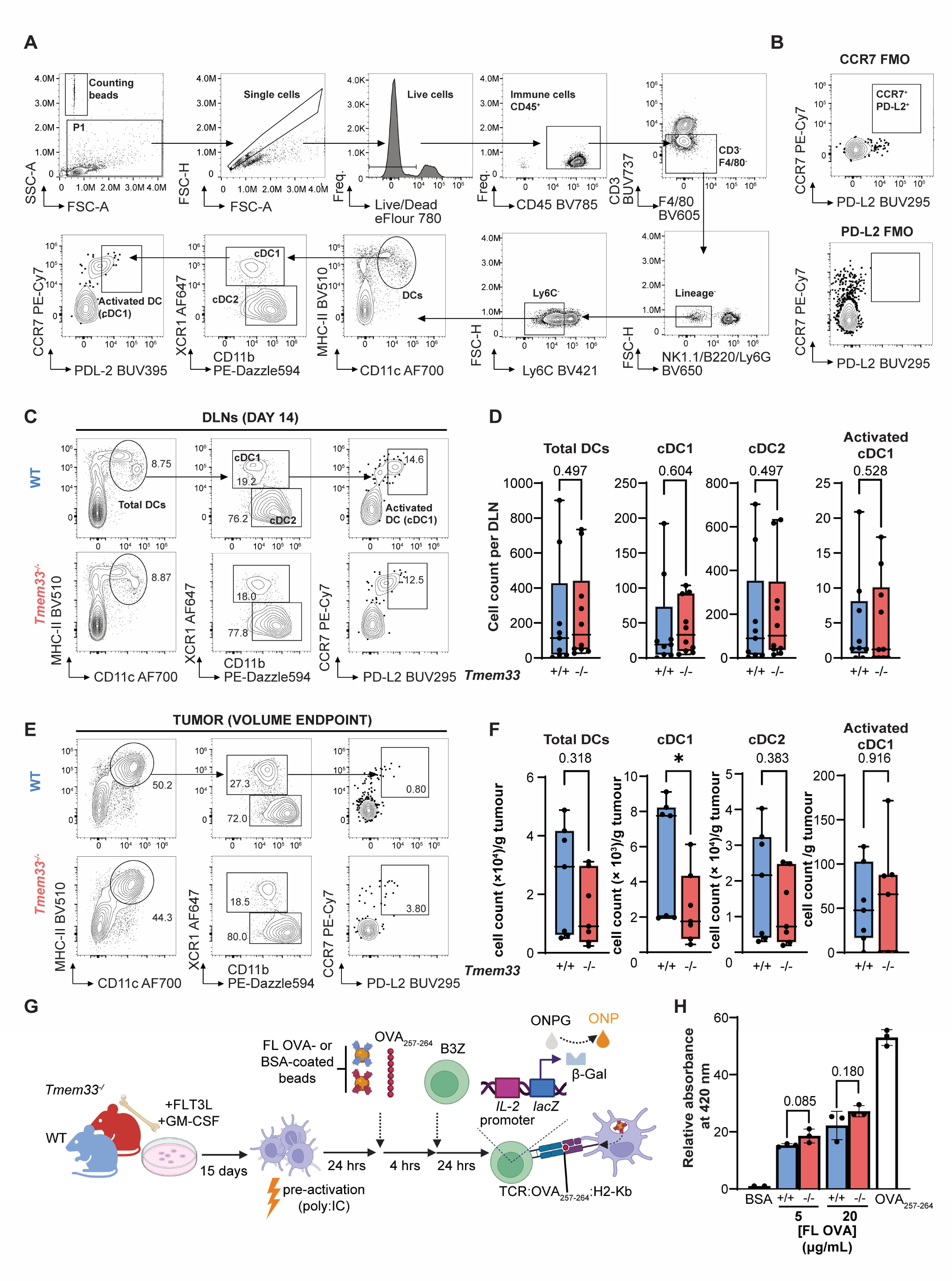
TMEM33 deletion did not alter dendritic cell (DC) abundance following tumor challenge, nor their cross-priming capacity *ex vivo*. **A**, Gating strategy employed for DC identification. DCs were defined as live, CD45^+^, lineage- (CD3, F4/80, NK1.1, B220, Ly6G, F4/80) and CD11c^+^MHCII^+^. cDC1, XCR1^+^CD11b^lo^; cDC2, XCR1^-^CD11b^hi^; activated DC, CCR7^+^PD- L2^+^. **B,** Fluorescence-minus-one (FMO) controls for CCR7 and PD-L2 markers. **C**, Representative plots for DC profiling in day 14 DLNs from B16F10-OVA tumor-bearing WT and *Tmem33^-/-^*female mice, with percentages of parent populations indicated. **D**, Absolute quantification of indicated DC subsets (depicted in **C**) from DLNs of individual mice. N = 9-10 mice/genotype. **E**, Representative plots (as in **C**) for DCs identified within B16F10-OVA tumors harvested upon reaching 600 mm^3^, and **F,** their absolute quantification in individual mice. N = 7 mice/genotype. **G**, Schematic depicting *ex vivo* examination of antigen cross-presentation in bone marrow-derived DCs (BMDCs). BMDCs derived using FLT3L and GM-CSF were pre-activated with polyI:C(HMW) for 24 hours before pulsing with BioMag®Plus Carboxyl beads (Polysciences) coated with full length (FL) ovalbumin (OVA), bovine serum albumin (BSA, negative control), or treating with OVA_257-264_ peptide (SIINFEKL, positive control) for 4 hours. Antigen-pulsed BMDCs were co-cultured with OVA_257-264_-restricted B3Z CD8^+^ T cell hybridomas for 24 hours, which upon activation upregulate β-galactosidase under control of the IL-2 promoter. β-galactosidase converts the colorimetric substrate ortho-Nitrophenyl-β-galactoside (ONPG) into ortho-Nitrophenyl (ONP). Designed using BioRender. **H**, Quantification of OVA_257-264_ cross-presentation in WT and *Tmem33^-/-^* BMDCs by monitoring B3Z-derived ONP absorbance at 420 nm. FL OVA beads were administered to BMDCs at both 5 and 20 µg/mL. N = 3 mice/genotype. DC counts in **D** and **F** were compared using Mann-Whitney *U* tests. Central lines and boxes indicate median and interquartile range; whiskers denote minimum and maximum values. In **H**, data were compared using unpaired two-tailed Student’s *t* tests. Bars and error represent mean and SD. \**, p≤*0.05; *p>*0.05 numerically indicated.

**Supplementary Figure 6.**
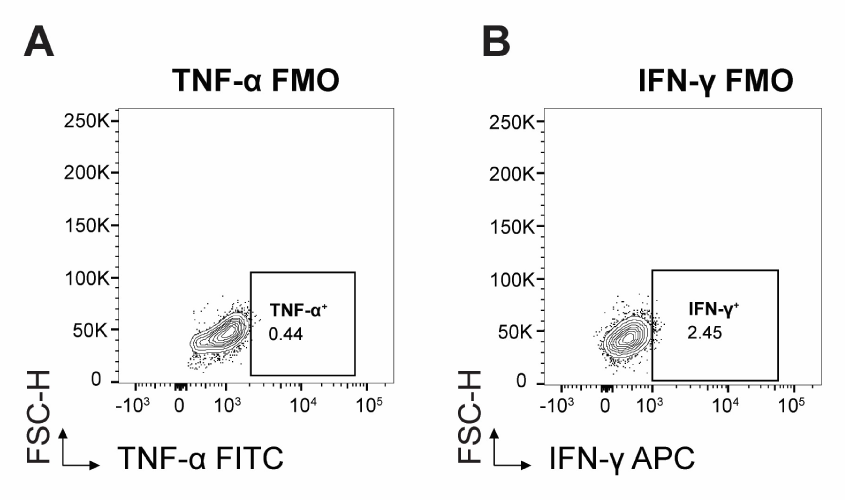
Supporting analyses for Figure 3: TMEM33 deficiency enhanced CD8^+^ T cell responses to polyclonal stimulation. **A**, **B,** Representative fluorescence-minus-one (FMO) gating controls used to define TNF-α^+^ (**A**) and IFN-γ^+^ (**B**) populations among naive CD8^+^ T cells following 24 hour stimulation with anti-CD3/CD28 Dynabeads® and recombinant IL-2.

**Supplementary Figure 7.**
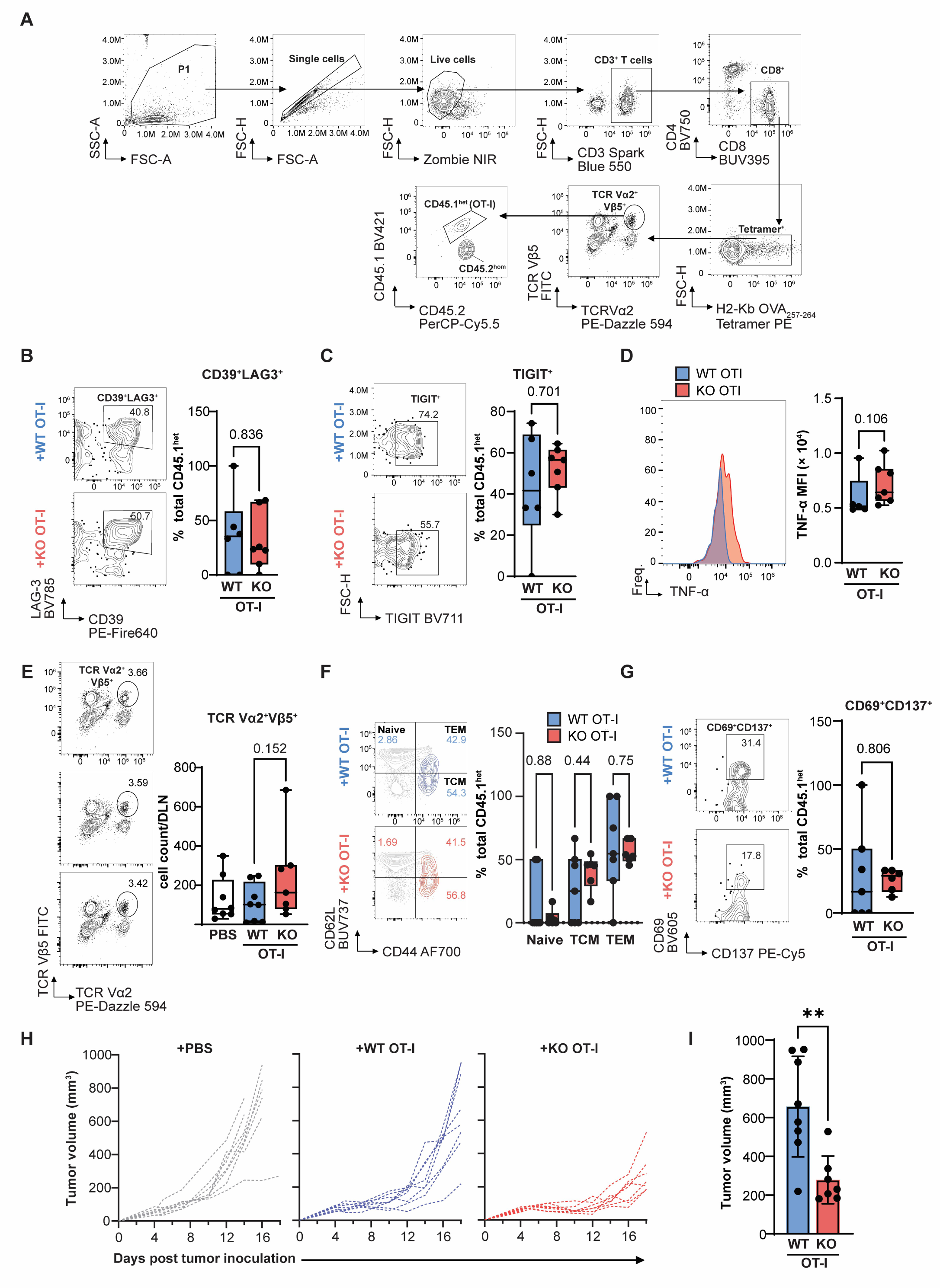
Gating strategy and additional flow cytometric profiling of transferred CD8^+^ OT-I T cells, and individual tumor growth profiling in mice receiving adoptive cell therapy. **A**, Full gating strategy employed to identify adoptively transferred CD45.1^het^ OT-I T cells. CD45.1^het^ cells were identified among tetramer^+^ populations co-expressing TCR Vα2 and TCR Vβ5 subunits. **B**, Representative plots (left) showing CD39^+^LAG3^+^ populations among *Tmem33^+/+^* (‘WT’) or *Tmem33^-/-^* (‘KO’) CD45.1^het^ OT-I cells in tumors (percentages indicated) and their quantification (right). **C,** Equivalent analysis (as in **B**) for TIGIT^+^ populations. **D,** TNF-α expression among indicated intratumoral OT-I populations with representative histograms for specified markers plotted (left), along with quantified MFI values for individual recipient mice (right). **E**, Representative plots indicating TCR Vα^2+^Vβ5^+^ populations among OVA_257-264_ tetramer^+^ CD8^+^ compartments in DLNs. Absolute cell counts were quantified in each recipient mouse (right). **F**, Characterization of naive (CD44^-^CD62L^+^), central memory (TCM; CD44^+^CD62L^+^) and effector memory (TEM; CD44^+^CD62L^-^) subsets among CD45.1^het^ OT-I cells in DLNs. In representative plots (left), transferred OT-I cells (blue or red) are superimposed on total CD8^+^ T cell populations (gray) in recipient mice. Percentages indicate subset frequencies among OT-I T cells and are quantified in adjacent box plots. **G**, Representative flow plots (left) and quantified frequencies (right) of CD45.1^het^ OT-I cells in DLNs co-expressing activation markers CD69 and CD137. N = 7-8 mice/treatment. Where no absolute CD45.1^het^ cells were identified (cell count = 0), these samples were excluded for downstream functional analysis. **H**, Growth profiles for B16F10-OVA tumors in individual mice receiving PBS, or pre-activated CD8^+^ T cells from WT (*Tmem33^+/+^*) OT-I or KO (*Tmem33^-/-^*) OT-I donor mice; **I,** Volumes were compared at day 18 post-implantation. Normality was assessed using Shapiro-Wilk tests. Non-parametric data are presented as box plots (central lines at median, boxes indicating interquartile range and whiskers showing range) and compared using Mann-Whitney *U* tests. For parametric data, bars and error represent mean and SD; data compared using unpaired two-tailed Student’s *t* tests. \**, p≤*0.05; \*\**p≤*0.01; *p>*0.05 numerically indicated.

**Supplementary Figure 8.**
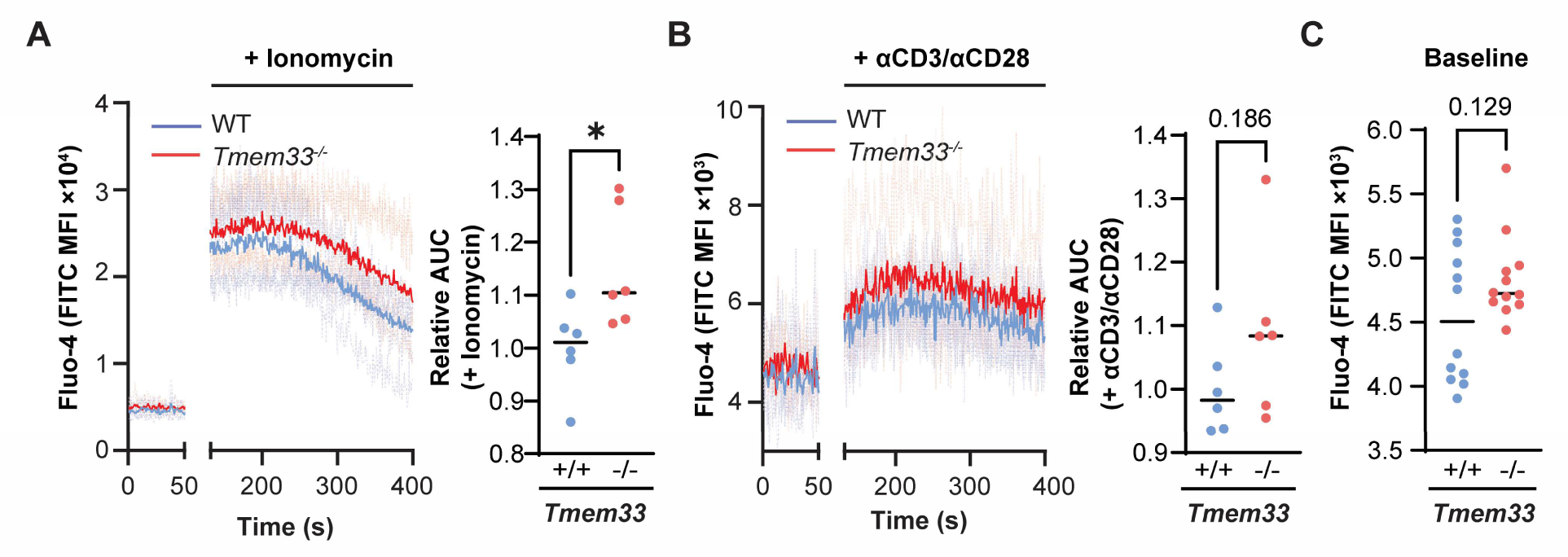
TMEM33 deletion increased Ca^2+^ mobilization in naive CD8^+^ T cells but did not affect TCR-induced transients. **A**, Intracellular Ca^2+^ monitoring by Fluo-4 staining in naïve CD8^+^ T cells in response to ionomycin treatment (10 µg/mL) or **B,** anti-CD3/CD28 antibody stimulation. Traces highlight Fluo-4 signal at baseline (0-50 seconds) and following stimulation (130-400 seconds). Solid lines, Fluo-4 MFI for each genotype; dashed lines, Fluo-4 MFI for individual samples. Relative area under curve (AUC) values (post-stimulation; 130-400 seconds) were generated by normalizing raw AUCs to the mean WT AUC within each experimental batch, correcting for inter-batch variability. **C,** Baseline Fluo-4 (FITC) mean fluorescence intensity (MFI) values for WT and *Tmem33^-/-^*naïve CD8^+^ T cells. N = 6 mice/genotype. For **C,** 2 technical replicates were recorded per mouse. All statistical comparisons were made using unpaired two-tailed Student’s *t* tests. \*\**, p≤*0.01; \**, p≤*0.05; *p>*0.05 numerically indicated. Bars represent mean.

**Supplementary Figure 9.**
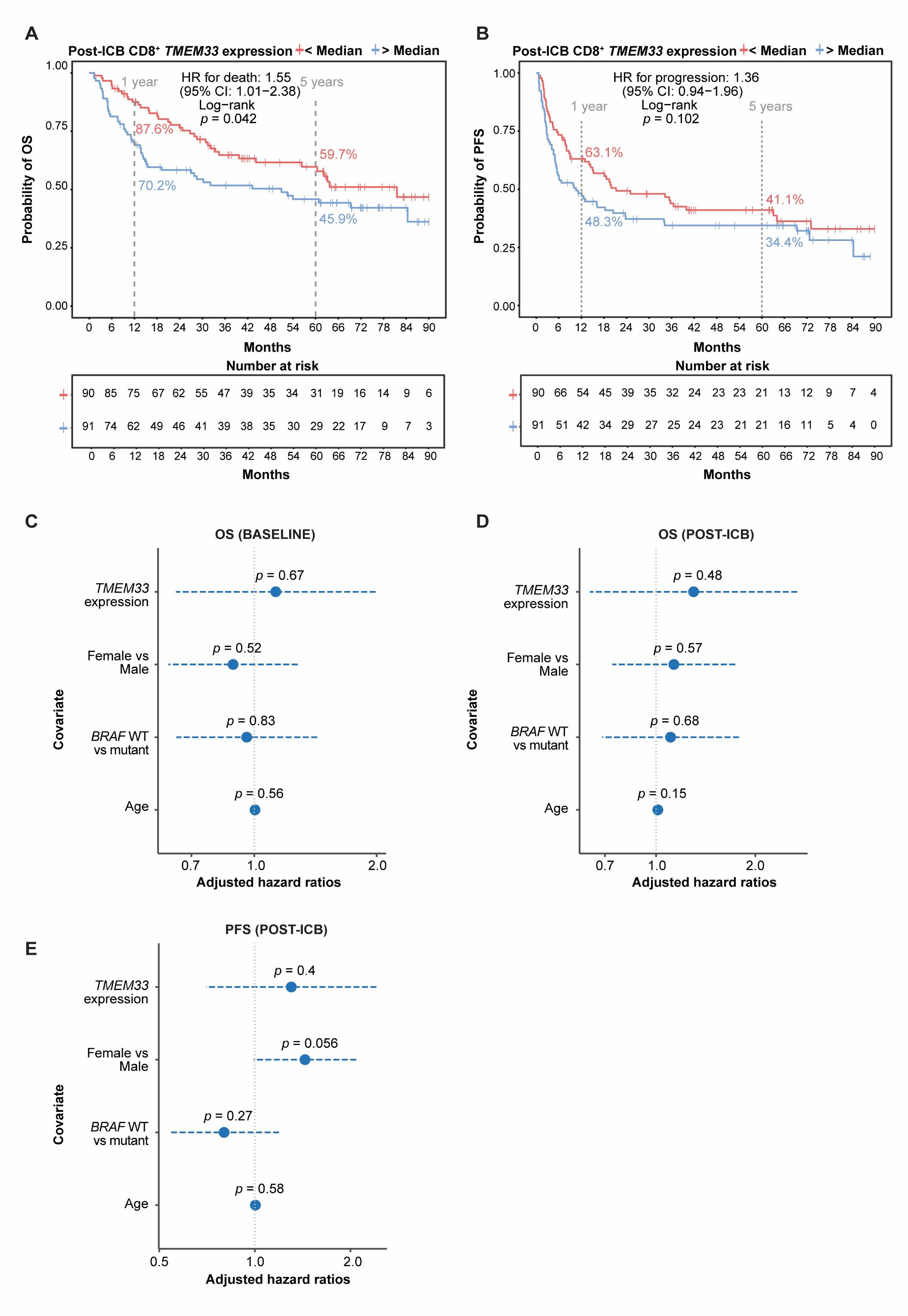
Supporting analyses for Figure 5: Reduced *TMEM33* expression in melanoma patient CD8^+^ T cells predicts improved survival and correlates with elevated *TCF7* expression. **A, B,** Kaplan-Meier analyses indicating overall survival (OS, **A**) and progression-free survival (PFS, **B**) in patients with below- or above-median *TMEM33* expression in peripheral CD8^+^ T cells determined 22 days following one cycle of immune checkpoint blockade (‘post-ICB’), by bulk RNA-seq. Two-sided log rank test. N below-median *TMEM33* = 90; N above-median *TMEM33* = 91. Hazard ratios (HR) for death (**A**) and progression (**B**) are indicated along with 95% confidence intervals (CI). **C,** Cox proportional hazards model examining the association between *TMEM33* expression and OS at baseline or **D**, post-ICB, adjusting for sex, age and *BRAF* status; Wald test. Dashed lines indicate CI. **E,** Equivalent analysis (as in **C** and **D**) for PFS post-ICB. For all analyses, *p* values are numerically indicated.

## REFERENCES

1. A. Cesano, R. Augustin, L. Barrea, D. Bedognetti, T. C. Bruno, A. Carturan, C. Hammer, W. S. Ho, J. N. Kather, T. Kirchhoff, R. O. Lu, J. McQuade, Y. G. Najjar, V. Pietrobon, M. Ruella, R. Shen, L. Soldati, C. Spencer, A. Betof Warner, S. Warren, E. Ziv, F. M. Marincola, Advances in the understanding and therapeutic manipulation of cancer immune responsiveness: a Society for Immunotherapy of Cancer (SITC) review. J Immunother Cancer 13 (2025).

2. D. R. Peaper, P. Cresswell, Regulation of MHC class I assembly and peptide binding. Annu Rev Cell Dev Biol 24, 343–68 (2008).

3. K. Shah, A. Al-Haidari, J. Sun, J. U. Kazi, T cell receptor (TCR) signaling in health and disease. Signal Transduct Target Ther 6, 412 (2021).

4. Y. Luo, L. Chang, Y. Ji, T. Liang, ER: a critical hub for STING signaling regulation. Trends Cell Biol 34, 865–881 (2024).

5. S.-M. Hwang, S. Chang, P. C. Rodriguez, J. R. Cubillos-Ruiz, Endoplasmic reticulum stress responses in anticancer immunity. Nat Rev Cancer 25, 684–702 (2025).

6. D. S. Schwarz, M. D. Blower, The endoplasmic reticulum: structure, function and response to cellular signaling. Cellular and Molecular Life Sciences 73, 79–94 (2016).

7. J. C. Christianson, P. Carvalho, Order through destruction: how ER-associated protein degradation contributes to organelle homeostasis. EMBO J 41, e109845 (2022).

8. E. J. Fenech, F. Lari, P. D. Charles, R. Fischer, M. Laétitia-Thézénas, K. Bagola, A. W. Paton, J. C. Paton, M. Gyrd-Hansen, B. M. Kessler, J. C. Christianson, Interaction mapping of endoplasmic reticulum ubiquitin ligases identifies modulators of innate immune signalling. Elife 9, e57306 (2020).

9. H. Chen, X. Zhao, Y. Li, S. Zhang, Y. Wang, L. Wang, W. Ma, High Expression of TMEM33 Predicts Poor Prognosis and Promotes Cell Proliferation in Cervical Cancer. Front Genet 13, 908807 (2022).

10. S. Loi, B. Haibe-Kains, C. Desmedt, F. Lallemand, A. M. Tutt, C. Gillet, P. Ellis, A. Harris, J. Bergh, J. A. Foekens, J. G. M. Klijn, D. Larsimont, M. Buyse, G. Bontempi, M. Delorenzi, M. J. Piccart, C. Sotiriou, Definition of Clinically Distinct Molecular Subtypes in Estrogen Receptor–Positive Breast Carcinomas Through Genomic Grade. Journal of Clinical Oncology 25, 1239–1246 (2007).

11. S. Loi, B. Haibe-Kains, S. Majjaj, F. Lallemand, V. Durbecq, D. Larsimont, A. M. Gonzalez-Angulo, L. Pusztai, W. F. Symmans, A. Bardelli, P. Ellis, A. N. J. Tutt, C. E. Gillett, B. T. Hennessy, G. B. Mills, W. A. Phillips, M. J. Piccart, T. P. Speed, G. A. McArthur, C. Sotiriou, PIK3CA mutations associated with gene signature of low mTORC1 signaling and better outcomes in estrogen receptor–positive breast cancer. Proceedings of the National Academy of Sciences 107, 10208–10213 (2010).

12. Y. Zhang, A. M. Sieuwerts, M. McGreevy, G. Casey, T. Cufer, A. Paradiso, N. Harbeck, P. N. Span, D. G. Hicks, J. Crowe, R. R. Tubbs, G. T. Budd, J. Lyons, F. C. G. J. Sweep, M. Schmitt, F. Schittulli, R. Golouh, D. Talantov, Y. Wang, J. A. Foekens, The 76-gene signature defines high-risk patients that benefit from adjuvant tamoxifen therapy. Breast Cancer Res Treat 116, 303–309 (2009).

13. I. Sakabe, R. Hu, L. Jin, R. Clarke, U. N. Kasid, TMEM33: a new stress-inducible endoplasmic reticulum transmembrane protein and modulator of the unfolded protein response signaling. Breast Cancer Res Treat 153, 285–97 (2015).

14. M. Arhatte, G. S. Gunaratne, C. El Boustany, I. Y. Kuo, C. Moro, F. Duprat, M. Plaisant, H. Duval, D. Li, N. Picard, A. Couvreux, C. Duranton, I. Rubera, S. Pagnotta, S. Lacas-Gervais, B. E. Ehrlich, J. S. Marchant, A. M. Savage, F. J. M. van Eeden, R. N. Wilkinson, S. Demolombe, E. Honoré, A. Patel, TMEM33 regulates intracellular calcium homeostasis in renal tubular epithelial cells. Nat Commun 10, 2024 (2019).

15. P.-L. Tsai, C. J. F. Cameron, M. F. Forni, R. R. Wasko, B. S. Naughton, V. Horsley, M. B. Gerstein, C. Schlieker, Dynamic quality control machinery that operates across compartmental borders mediates the degradation of mammalian nuclear membrane proteins. Cell Rep 41, 111675 (2022).

16. F. Liu, M. Ma, A. Gao, F. Ma, G. Ma, P. Liu, C. Jia, Y. Wang, K. Donahue, S. Zhang, I. M. Ong, S. Keles, L. Li, W. Xu, PKM2-TMEM33 axis regulates lipid homeostasis in cancer cells by controlling SCAP stability. EMBO J 40, e108065 (2021).

17. D. T. Utzschneider, L. Charmoy, V. Chennupati, R. Thimme, D. Zehn, W. Held, T Cell Factor 1-Expressing Memory-like CD8 + T Cells Sustain the Immune Response to Chronic Viral Infections. Immunity 45, 415–427 (2016).

18. M. Sade-Feldman, K. Yizhak, S. L. Bjorgaard, J. P. Ray, C. G. de Boer, R. W. Jenkins, D. J. Lieb, J. H. Chen, D. T. Frederick, M. Barzily-Rokni, S. S. Freeman, A. Reuben, P. J. Hoover, A.-C. Villani, E. Ivanova, A. Portell, P. H. Lizotte, A. R. Aref, J.-P. Eliane, M. R. Hammond, H. Vitzthum, S. M. Blackmon, B. Li, V. Gopalakrishnan, S. M. Reddy, Z. A. Cooper, C. P. Paweletz, D. A. Barbie, A. Stemmer-Rachamimov, K. T. Flaherty, J. A. Wargo, G. M. Boland, R. J. Sullivan, G. Getz, N. Hacohen, Defining T Cell States Associated with Response to Checkpoint Immunotherapy in Melanoma. Cell 175, 998–1013.e20 (2018).

19. B. C. Miller, D. R. Sen, R. Al Abosy, K. Bi, Y. V Virkud, M. W. LaFleur, K. B. Yates, A. Lako, K. Felt, G. S. Naik, M. Manos, E. Gjini, J. R. Kuchroo, J. J. Ishizuka, J. L. Collier, G. K. Griffin, S. Maleri, D. E. Comstock, S. A. Weiss, F. D. Brown, A. Panda, M. D. Zimmer, R. T. Manguso, F. S. Hodi, S. J. Rodig, A. H. Sharpe, W. N. Haining, Subsets of exhausted CD8+ T cells differentially mediate tumor control and respond to checkpoint blockade. Nat Immunol 20, 326–336 (2019).

20. I. Siddiqui, K. Schaeuble, V. Chennupati, S. A. Fuertes Marraco, S. Calderon-Copete, D. Pais Ferreira, S. J. Carmona, L. Scarpellino, D. Gfeller, S. Pradervand, S. A. Luther, D. E. Speiser, W. Held, Intratumoral Tcf1 + PD-1 + CD8 + T Cells with Stem-like Properties Promote Tumor Control in Response to Vaccination and Checkpoint Blockade Immunotherapy. Immunity 50, 195–211.e10 (2019).

21. S. J. Im, B. T. Konieczny, W. H. Hudson, D. Masopust, R. Ahmed, PD-1+ stemlike CD8 T cells are resident in lymphoid tissues during persistent LCMV infection. Proc Natl Acad Sci U S A 117, 4292–4299 (2020).

22. J. Brummelman, E. M. Mazza, G. Alvisi, F. S. Colombo, A. Grilli, J. Mikulak, D. Mavilio, M. Alloisio, F. Ferrari, E. Lopci, P. Novellis, G. Veronesi, E. Lugli, High-dimensional single cell analysis identifies stem-like cytotoxic CD8 + T cells infiltrating human tumors. Journal of Experimental Medicine 215, 2520–2535 (2018).

23. L.-F. Lu, C. Zhang, Z.-C. Li, X.-Y. Zhou, J.-Y. Jiang, D.-D. Chen, Y.-A. Zhang, F. Xiong, F. Zhou, S. Li, A novel role of Zebrafish TMEM33 in negative regulation of interferon production by two distinct mechanisms. PLoS Pathog 17, e1009317 (2021).

24. M. Shen, X. Jiang, Q. Peng, L. Oyang, Z. Ren, J. Wang, M. Peng, Y. Zhou, X. Deng, Q. Liao, The cGAS‒STING pathway in cancer immunity: mechanisms, challenges, and therapeutic implications. J Hematol Oncol 18, 40 (2025).

25. B. M. Sullivan, A. Juedes, S. J. Szabo, M. von Herrath, L. H. Glimcher, Antigen-driven effector CD8 T cell function regulated by T-bet. Proc Natl Acad Sci U S A 100, 15818–23 (2003).

26. S. Kurtulus, A. Madi, G. Escobar, M. Klapholz, J. Nyman, E. Christian, M. Pawlak, D. Dionne, J. Xia, O. Rozenblatt-Rosen, V. K. Kuchroo, A. Regev, A. C. Anderson, Checkpoint Blockade Immunotherapy Induces Dynamic Changes in PD-1-CD8+ Tumor-Infiltrating T Cells. Immunity 50, 181–194.e6 (2019).

27. M. E. Mikucki, D. T. Fisher, J. Matsuzaki, J. J. Skitzki, N. B. Gaulin, J. B. Muhitch, A. W. Ku, J. G. Frelinger, K. Odunsi, T. F. Gajewski, A. D. Luster, S. S. Evans, Non-redundant requirement for CXCR3 signalling during tumoricidal T-cell trafficking across tumour vascular checkpoints. Nat Commun 6, 7458 (2015).

28. N. Karin, Chemokines and cancer: new immune checkpoints for cancer therapy. Curr Opin Immunol 51, 140–145 (2018).

29. M. T. Chow, A. J. Ozga, R. L. Servis, D. T. Frederick, J. A. Lo, D. E. Fisher, G. J. Freeman, G. M. Boland, A. D. Luster, Intratumoral Activity of the CXCR3 Chemokine System Is Required for the Efficacy of Anti-PD-1 Therapy. Immunity 50, 1498–1512.e5 (2019).

30. N. Karin, CXCR3 Ligands in Cancer and Autoimmunity, Chemoattraction of Effector T Cells, and Beyond. Front Immunol 11, 976 (2020).

31. I. F. Hermans, D. S. Ritchie, J. Yang, J. M. Roberts, F. Ronchese, CD8+ T Cell-Dependent Elimination of Dendritic Cells In Vivo Limits the Induction of Antitumor Immunity. The Journal of Immunology 164, 3095–3101 (2000).

32. M. Chen, Y.-H. Wang, Y. Wang, L. Huang, H. Sandoval, Y.-J. Liu, J. Wang, Dendritic Cell Apoptosis in the Maintenance of Immune Tolerance. Science (1979) 311, 1160–1164 (2006).

33. S. Sanderson, N. Shastri, LacZ inducible, antigen/MHC-specific T cell hybrids. Int Immunol 6, 369–76 (1994).

34. K. A. Hogquist, S. C. Jameson, W. R. Heath, J. L. Howard, M. J. Bevan, F. R. Carbone, T cell receptor antagonist peptides induce positive selection. Cell 76, 17–27 (1994).

35. A. M. Savage, S. Kurusamy, Y. Chen, Z. Jiang, K. Chhabria, R. B. MacDonald, H. R. Kim, H. L. Wilson, F. J. M. van Eeden, A. L. Armesilla, T. J. A. Chico, R. N. Wilkinson, tmem33 is essential for VEGF-mediated endothelial calcium oscillations and angiogenesis. Nat Commun 10, 732 (2019).

36. G. Milotay, M. Little, R. A. Watson, D. Muldoon, S. MacKay, A. Kurioka, O. Tong, C. A. Taylor, I. Nassiri, L. M. Webb, O. Akin-Adigun, J. Bremke, W. Ye, B. Sun, P. K. Sharma, R. Cooper, S. Danielli, F. M. Santo, A. Verge de Los Aires, G. Niu, L. Cohen, E. Ng, J. J. Gilchrist, A. Y. Chong, A. Mentzer, V. Woodcock, N. Coupe, M. J. Payne, M. Youdell, M. R. Middleton, P. Klenerman, B. P. Fairfax, CMV serostatus is associated with improved survival and delayed toxicity onset following anti-PD-1 checkpoint blockade. Nat Med 31, 2350–2364 (2025).

37. B. P. Fairfax, C. A. Taylor, R. A. Watson, I. Nassiri, S. Danielli, H. Fang, E. A. Mahé, R. Cooper, V. Woodcock, Z. Traill, M. H. Al-Mossawi, J. C. Knight, P. Klenerman, M. Payne, M. R. Middleton, Peripheral CD8+ T cell characteristics associated with durable responses to immune checkpoint blockade in patients with metastatic melanoma. Nat Med 26, 193–199 (2020).

38. N. Prokhnevska, M. A. Cardenas, R. M. Valanparambil, E. Sobierajska, B. G. Barwick, C. Jansen, A. Reyes Moon, P. Gregorova, L. delBalzo, R. Greenwald, M. A. Bilen, M. Alemozaffar, S. Joshi, C. Cimmino, C. Larsen, V. Master, M. Sanda, H. Kissick, CD8+ T cell activation in cancer comprises an initial activation phase in lymph nodes followed by effector differentiation within the tumor. Immunity 56, 107–124.e5 (2023).

39. Z. Li, Z. K. Tuong, I. Dean, C. Willis, F. Gaspal, R. Fiancette, S. Idris, B. Kennedy, J. R. Ferdinand, A. Peñalver, M. Cabantous, S. M. Baker, J. W. Fry, G. Carlesso, S. A. Hammond, S. J. Dovedi, M. R. Hepworth, M. R. Clatworthy, D. R. Withers, In vivo labeling reveals continuous trafficking of TCF-1+ T cells between tumor and lymphoid tissue. Journal of Experimental Medicine 219, e20210749 (2022).

40. Y. Ma, Y. Wang, X. Zhao, G. Jin, J. Xu, Z. Li, N. Yin, Z. Gao, B. Xia, M. Peng, TMEM41B is an endoplasmic reticulum Ca2+ release channel maintaining naive T cell quiescence and responsiveness. Cell Discov 11, 18 (2025).

41. M. Zheng, W. Zhang, X. Chen, H. Guo, H. Wu, Y. Xu, Q. He, L. Ding, B. Yang, The impact of lipids on the cancer-immunity cycle and strategies for modulating lipid metabolism to improve cancer immunotherapy. Acta Pharm Sin B 13, 1488–1497 (2023).

42. Y. Li, R. Tinoco, L. Elmén, I. Segota, Y. Xian, Y. Fujita, A. Sahu, R. Zarecki, K. Marie, Y. Feng, A. Khateb, D. T. Frederick, S. K. Ashkenazi, H. Kim, E. G. Perez, C.-P. Day, R. S. Segura Muñoz, R. Schmaltz, S. Yooseph, M. A. Tam, T. Zhang, E. Avitan-Hersh, L. Tzur, S. Roizman, I. Boyango, G. Bar-Sela, A. Orian, R. J. Kaufman, M. Bosenberg, C. R. Goding, B. Baaten, M. P. Levesque, R. Dummer, K. Brown, G. Merlino, E. Ruppin, K. Flaherty, A. Ramer-Tait, T. Long, S. N. Peterson, L. M. Bradley, Z. A. Ronai, Gut microbiota dependent anti-tumor immunity restricts melanoma growth in Rnf5-/- mice. Nat Commun 10, 1492 (2019).

43. R. C. Sterner, R. M. Sterner, CAR-T cell therapy: current limitations and potential strategies. Blood Cancer J 11, 69 (2021).

44. A. Schietinger, M. Philip, V. E. Krisnawan, E. Y. Chiu, J. J. Delrow, R. S. Basom, P. Lauer, D. G. Brockstedt, S. E. Knoblaugh, G. J. Hämmerling, T. D. Schell, N. Garbi, P. D. Greenberg, Tumor-Specific T Cell Dysfunction Is a Dynamic Antigen-Driven Differentiation Program Initiated Early during Tumorigenesis. Immunity 45, 389–401 (2016).

45. B. R. Ford, P. D. A. Vignali, N. L. Rittenhouse, N. E. Scharping, R. Peralta, K. Lontos, A. T. Frisch, G. M. Delgoffe, A. C. Poholek, Tumor microenvironmental signals reshape chromatin landscapes to limit the functional potential of exhausted T cells. Sci Immunol 7, eabj9123 (2022).

46. A. T. Frisch, Y. Wang, B. Xie, A. Yang, B. R. Ford, S. Joshi, K. M. Kedziora, R. Peralta, D. Wilfahrt, S. J. Mullett, K. Spahr, K. Lontos, J. A. Jana, V. G. Dean, W. G. Gunn, S. Gelhaus, A. C. Poholek, D. B. Rivadeneira, G. M. Delgoffe, Redirecting glucose flux during in vitro expansion generates epigenetically and metabolically superior T cells for cancer immunotherapy. Cell Metab 37, 870–885.e8 (2025).

47. I. Pallavicini, T. M. Frasconi, C. Catozzi, E. Ceccacci, S. Tiberti, D. Haas, J. Samson, C. Heuser-Loy, C. B. Nava Lauson, M. Mangione, E. Preto, A. Bigogno, E. Sala, M. Iannacone, C. Mercurio, L. Gattinoni, I. Caruana, M. Kuka, L. Nezi, S. Minucci, T. Manzo, LSD1 inhibition improves efficacy of adoptive T cell therapy by enhancing CD8+ T cell responsiveness. Nat Commun 15, 7366 (2024).

48. R. P. Patel, G. Ghilardi, Y. Zhang, Y.-H. Chiang, W. Xie, P. Guruprasad, K. H. Kim, I. Chun, M. G. Angelos, R. Pajarillo, S. J. Hong, Y. G. Lee, O. Shestova, C. Shaw, I. Cohen, A. Gupta, T. Vu, D. Qian, S. Yang, A. Nimmagadda, A. E. Snook, N. Siciliano, A. Rotolo, A. Inamdar, J. Harris, O. Ugwuanyi, M. Wang, A. Carturan, L. Paruzzo, L. Chen, H. J. Ballard, T. Blanchard, C. Xu, M. Abdel-Mohsen, K. Gabunia, M. Wysocka, G. P. Linette, B. Carreno, D. M. Barrett, D. T. Teachey, A. D. Posey, D. J. Powell, C. T. Sauter, S. Pileri, V. Pillai, J. Scholler, A. H. Rook, S. J. Schuster, S. K. Barta, P. Porazzi, M. Ruella, CD5 deletion enhances the antitumor activity of adoptive T cell therapies. Sci Immunol 9, eadn6509 (2024).

49. V. De Castro, J. Galaine, R. Loyon, Y. Godet, CRISPR-Cas gene knockouts to optimize engineered T cells for cancer immunotherapy. Cancer Gene Ther 31, 1124–1134 (2024).

50. L. Jin, K. K. Hill, H. Filak, J. Mogan, H. Knowles, B. Zhang, A.-L. Perraud, J. C. Cambier, L. L. Lenz, MPYS is required for IFN response factor 3 activation and type I IFN production in the response of cultured phagocytes to bacterial second messengers cyclic-di-AMP and cyclic-di-GMP. J Immunol 187, 2595–601 (2011).

51. T. M. Ashhurst, F. Marsh-Wakefield, G. H. Putri, A. G. Spiteri, D. Shinko, M. N. Read, A. L. Smith, N. J. C. King, Integration, exploration, and analysis of high-dimensional single-cell cytometry data using Spectre. Cytometry Part A 101, 237–253 (2022).

52. M. Martin, Cutadapt removes adapter sequences from high-throughput sequencing reads. EMBnet J 17, 10 (2011).

53. A. Dobin, C. A. Davis, F. Schlesinger, J. Drenkow, C. Zaleski, S. Jha, P. Batut, M. Chaisson, T. R. Gingeras, STAR: ultrafast universal RNA-seq aligner. Bioinformatics 29, 15–21 (2013).

54. Y. Liao, G. K. Smyth, W. Shi, featureCounts: an efficient general purpose program for assigning sequence reads to genomic features. Bioinformatics 30, 923–30 (2014).

55. M. I. Love, W. Huber, S. Anders, Moderated estimation of fold change and dispersion for RNA-seq data with DESeq2. Genome Biol 15, 550 (2014).

56. A. Subramanian, P. Tamayo, V. K. Mootha, S. Mukherjee, B. L. Ebert, M. A. Gillette, A. Paulovich, S. L. Pomeroy, T. R. Golub, E. S. Lander, J. P. Mesirov, Gene set enrichment analysis: A knowledge-based approach for interpreting genome-wide expression profiles. Proceedings of the National Academy of Sciences 102, 15545–15550 (2005).

57. G. Korotkevich, V. Sukhov, N. Budin, B. Shpak, M. N. Artyomov, A. Sergushichev, Fast gene set enrichment analysis. bioRxiv 060012 (2016).

58. A. Liberzon, A. Subramanian, R. Pinchback, H. Thorvaldsdóttir, P. Tamayo, J. P. Mesirov, Molecular signatures database (MSigDB) 3.0. Bioinformatics 27, 1739–40 (2011).

59. M. Ashburner, C. A. Ball, J. A. Blake, D. Botstein, H. Butler, J. M. Cherry, A. P. Davis, K. Dolinski, S. S. Dwight, J. T. Eppig, M. A. Harris, D. P. Hill, L. Issel-Tarver, A. Kasarskis, S. Lewis, J. C. Matese, J. E. Richardson, M. Ringwald, G. M. Rubin, G. Sherlock, Gene Ontology: tool for the unification of biology. Nat Genet 25, 25–29 (2000).

60. M. Schemper, T. L. Smith, A note on quantifying follow-up in studies of failure time. Control Clin Trials 17, 343–6 (1996).

